# Protein dynamics affect O_2_-stability of Group B [FeFe]-hydrogenase from *Thermosediminibacter oceani*

**DOI:** 10.1101/2025.03.17.643706

**Authors:** Subhasri Ghosh, Chandan K. Das, Sarmila Uddin, Sven T. Stripp, Vera Engelbrecht, Martin Winkler, Silke Leimkühler, Claudia Brocks, Jifu Duan, Lars V. Schäfer, Thomas Happe

**Affiliations:** Photobiotechnology, Ruhr University Bochum, 44801 Bochum, Germany; Center for Theoretical Chemistry, Ruhr University Bochum, 44801 Bochum, Germany; Computational Biotechnology, RWTH Aachen University, 52074 Aachen, Germany; Technische Universität Berlin, Division of Physical Chemistry, Strasse des 17. Juni 124, 10623 Berlin, Germany; Electrobiotechnology, Technical University of Munich Campus Straubing for Biotechnology and Sustainability, 94315 Straubing, Germany; Molecular Enzymology, University of Potsdam, Karl-Liebknecht-Str. 24, 14476 Potsdam, Germany

**Keywords:** [FeFe]-hydrogenases, O_2_-stability, Hydrophobic cluster, IR spectroscopy, MD simulations

## Abstract

In the pursuit of sustainable ‘green’ energy generation, [FeFe]-hydrogenases have attracted significant attention due to their ability to catalyze hydrogen production. However, the sensitivity of these enzymes to O_2_ is a major obstacle for their application as biocatalysts in energy conversion technologies. In the search for an O_2_-stable [FeFe]-hydrogenase, we identified the hydrogenase ToHydA from *Thermosediminibacter oceani* that belongs to the rarely characterized Group B (M2a) [FeFe]-hydrogenases. Our findings demonstrate that ToHydA exhibits remarkable O_2_-stability, even under prolonged O_2_ exposure. By characterizing site-directed mutagenesis variants, we found that the highly conserved proton-transporting cysteine protects H-cluster from O_2_-induced degradation by forming H_inact_ state. The additional cysteine residue in the TSCCCP motif of ToHydA, a feature unique to Group B (M2a) [FeFe]-hydrogenases, enhances the flexibility of that motif and facilitates the formation of the H_inact_ state. Moreover, ToHydA possesses unique features, including the formation of an unusual H_inact_ resting state that distinguishes the enzyme from other [FeFe]-hydrogenases. Our atomistic molecular dynamics simulations reveal a previously unrecognized cluster of hydrophobic residues centered around the proton-transporting cysteine-bearing loop. This structural feature appears to be a common molecular characteristic in hydrogenases that form the O_2_-protected H_inact_ state. By exploiting these molecular features of ToHydA, future research can aim to rationally design hydrogenases that combine high catalytic activity with enhanced O_2_ stability, to develop more efficient and durable catalysts.

**Table of Contents Figure:** 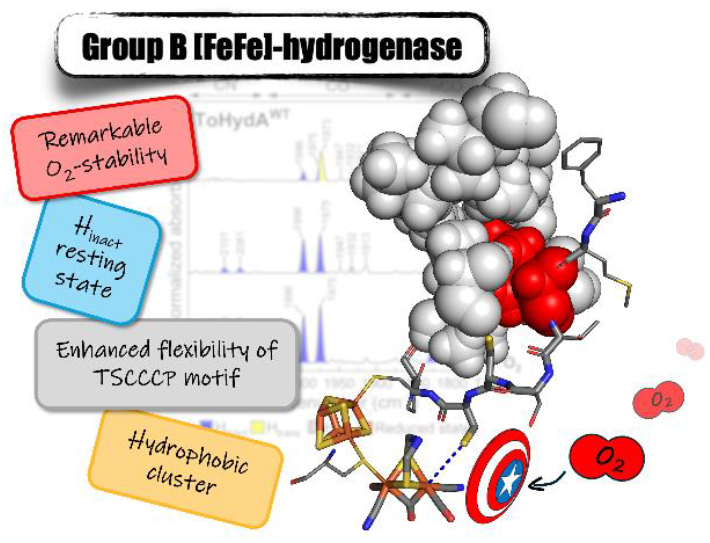

## Introduction

Molecular hydrogen (H_2_) is a promising ‘clean’ alternative to fossil fuels due to its combustion, which yields only water without producing greenhouse gases, thus mitigating global warming.^1,2^ In contrast to industrial H_2_ production, which primarily relies on steam reforming of natural gas,^3^ nature employs highly efficient catalysts known as [FeFe]-hydrogenases. Phylogenetic analysis of [FeFe]-hydrogenase protein sequences reveals primarily the existence of four main groups, labeled A-D, each containing several subclasses.^4–7^ Group A hydrogenases are prototypical and bifurcating, involved in both H_2_ oxidation and production. Well-studied examples include CrHydA1 from *Chlamydomonas reinhardtii*,^8–10^ CpI from *Clostridium pasteurianum*,^11,12^ and DdH from *Desulfovibrio desulfuricans*.^13–15^ Groups C and D comprise sensory-type hydrogenases.^16–19^ The Group B hydrogenases remains mostly unexplored; however, they are considered ancestral types of [FeFe]-hydrogenases and recently gained significant attention from the scientific community.^20–22^ The current study focuses mainly on Group B (M2a subgroup) [FeFe]-hydrogenases. Recently, classification of hydrogenases is extended by the addition of groups E, F and G that comprise of ultraminimal [FeFe]-hydrogenases and and hybrid hydrogenases.^23^

The active site of [FeFe]-hydrogenase, termed the H-cluster, is deeply embedded within the enzyme’s protein matrix. It consists of a canonical [4Fe-4s]-cluster ([4Fe]_H_) and a unique diiron site ([2Fe]_H_) connected via a bridging cysteine residue.^11,13,24–26^ The [2Fe]_H_-subsite is coordinated by carbon monoxide (CO) and cyanide (CN−) ligands, which stabilize the low oxidation states of the diiron ions of the [2Fe]_H_. The two iron atoms in the [2Fe]_H_-subsite are termed as proximal Fe (Fe_p_) and distal Fe (Fe_d_), according to their position relative to the [4Fe]_H_-cluster. The [2Fe]_H_-subsite is further coordinated by an azadithiolate (adt) bridge, where the nitrogen atom serves as a proton receptor for catalysis. The Fe_d_ site is pentacoordinated, with its open coordination site capable of reversibly binding to H^+^ or H_2_, depending on applied conditions. This site can be inactivated by binding to other ligands (CO, CN^-^, HS^-^, O_2_, etc.).^27–35^

The industrial application of [FeFe]-hydrogenases is significantly impeded by their susceptibility to irreversible inactivation upon exposure to O_2_, which leads to the degradation of their active sites.^36^ Although the exact mechanism of O_2_-induced inactivation remains to be fully elucidated through direct experimental evidence, computational studies propose a plausible pathway.^37,38^ In this proposed degradation mechanism, O_2_ binds to the vacant coordination site of Fe_d_ and forms a superoxide ion. Subsequent electron transfer and protonation steps generate hydroperoxyl radical and hydrogen peroxide, collectively termed as reactive oxygen species (ROS). These reactive species subsequently attack and degrade both the [2Fe]_H_-cluster and the [4Fe]_H_-cubane.^34–36,39,40^ Recent studies have shown that in CpI, the diffusion of O_2_ or ROS between the [2Fe]_H_-cluster and [4Fe]_H_-cluster occurs through a water pathway which is a crucial step for H-cluster degradation. Point mutations that block this water pathway, can significantly reduce the diffusion of O_2_ or ROS, thereby leading to more O_2_-tolerant hydrogenase.^41^ However, engineering fully O_2_-tolerant hydrogenase by mutagenesis to impede such ROS diffusion is challenging.

An alternative mechanism of O_2_ protection has been reported recently in CbA5H, a Group A [FeFe]-hydrogenase from *Clostridium beijerinckii*.^47^ The CbA5H exhibits an unique mechanism of O_2_ protection by reversibly switching between the catalytically active oxidized state (H_ox_) and the O_2_-protected H_inact_ state.^44,48–50^ In the presence of O_2_, the thiol (-SH) group of the proton-transporting cysteine (C367 in CbA5H),^51^ coordinates to the Fe_d_ of CbA5H, thereby preventing O_2_ binding and subsequent damage to the active site.^44^ Similar to CbA5H, Artz et al. identified oxidative inactivation in CpIII, a Group B (M2a) [FeFe]-hydrogenase from *Clostridium pasteurianum*, despite their classification into different phylogenetic groups within the [FeFe]-hydrogenase family.^20,21^ Group B (M2a) hydrogenases have an additional cysteine residue next to the proton transporting cysteine residue, forming a TSCCCP motif instead of the highly conserved TSCCP motif of Group A hydrogenases; see Figure 1 showing a comparative sequence alignment between Group A and B [FeFe]-hydrogenases (sequence alignment of 144 [FeFe]-hydrogenases is shown in SI Figure S1). Detailed electrochemical characterization by Fasano et al. confirms that CpIII undergoes oxidative inactivation through the formation of two inactive species. The authors hypothesized that in one of the species, the proton-transporting cysteine binds to Fe_d_. However, the exact identity of these inactive species remains unclear due to the lack of spectroscopic evidence._22_

**Figure 1.**
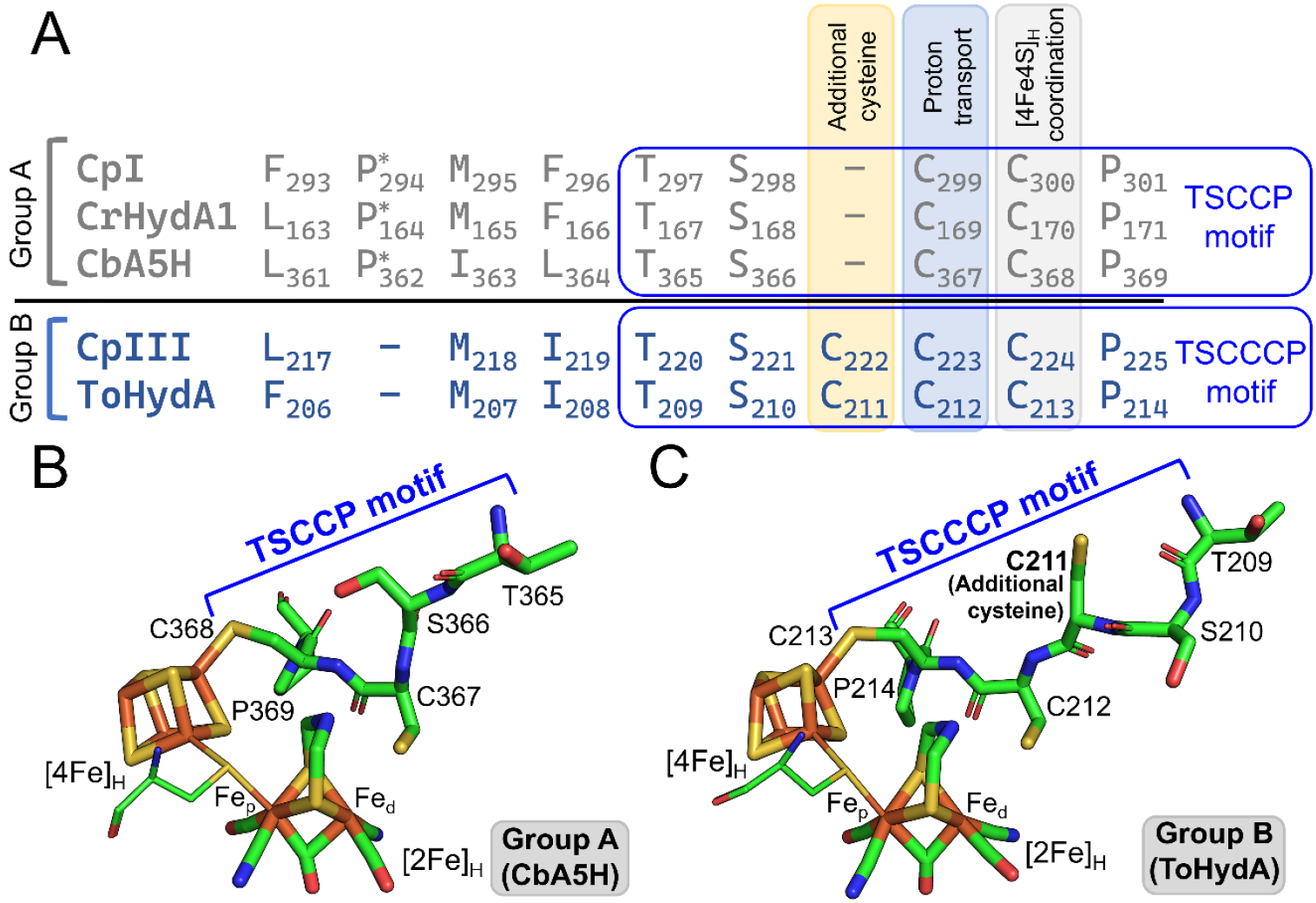
**A.** Sequence alignment of the proton-transporting cysteine-bearing loop from Group A [FeFe]-hydrogenases (CpI, CrHydA1, and CbA5H) and Group B (M2a) [FeFe]-hydrogenases (CpIII and ToHydA). This is part of the sequence alignment of 144 [FeFe]-hydrogenases from HydDB^42^ (72 Group B (M2a) with TSCCCP motif and 72 Group A with TSCCP motif) generated using Clustal Omega^43^ (pdf file containing the sequence alignment is provided as supporting information). For details refer to SI Figure S1. The additional cysteine residue for Group B (M2a), the proton transporting and [4Fe]_H_-coordinating cysteine residue for Group A and B [FeFe]-hydrogenase are also indicated in the figure. The asterisk marks the proline residues that are present in Group A [FeFe]-hydrogenases but absent in Group B (M2a). B and C. Structural views of the active sites of Group A (CbA5H, PDB: 6TTL)^44^ and Group B (M2a) (ToHydA, generated using AlphaFold,^45,46^ details in SI MD details) [FeFe]-hydrogenases are shown, respectively. The TSCCP motif for Group A, as well as the TSCCCP motif along with additional cysteine residue for Group B (M2a), are highlighted in the figures.

A manually curated hydrogenase database (HydDB) was established and updated by Søndergaard et al. to classify all the hydrogenases from different organisms.^42^ We performed a thorough database search in HydDB and the NCBI database to identify other Group B (M2a) hydrogenases. A [FeFe]-hydrogenase from *Thermosediminibacter oceani* (hereafter referred to as ToHydA), an anaerobic thermophilic bacterium, is identified and thoroughly characterized in this study. Notably, the ToHydA exhibits typical protein sequence features of Group B (M2a) [FeFe]-hydrogenases, particularly the presence of an additional cysteine residue adjacent to the proton-transporting cysteine, forming the TSCCCP motif. Through a series of biochemical and spectroscopic experiments, we characterized the wild-type (WT) ToHydA enzyme and several of its variants, assessing their abilities to protect H-cluster against O_2_ induced degradation. Interestingly, our spectroscopic study also identified a non-canonical resting state for the WT enzyme and a potential secondary inactive state for the variants that do not form the H_inact_ state in the presence of O_2_. Additionally, our all-atom molecular dynamics (MD) simulations revealed the dynamic network of backbone hydrogen bonds between the TSCCCP motif and surrounding residues, as well as previously unrecognized hydrophobic cluster formations playing pivotal roles in the H_inact_ formation. These crucial molecular determinants modulating the dynamics of the TSCCCP motif during H_inact_ formation pave the way for engineering O_2_-stable [FeFe]-hydrogenases.

## Results

### 2.1. Recombinant ToHydA WT is an active [FeFe]-hydro-genase

The apo-protein of ToHydA (47.3 kDa), lacking the [2Fe]_H_ cluster, was heterologously expressed in *E. coli* without the maturases under anaerobic conditions, followed by purification using affinity chromatography (detailed in the methods section, see SI; SDS-PAGE shown in SI Figure S2). Subsequently, the isolated protein was maturated with synthetic [2Fe]^MIM^ to produce the holo-protein (hereafter referred to as ToHydA WT) used for further experiments. The iron content (Fe atoms per protein molecule) of ToHydA WT was determined using inductively coupled plasma-optical emission spectroscopy (ICP-OES) for both the apo and holo forms, revealing (10.76 ± 0.18) and (13.29 ± 0.04) iron atoms, respectively (SI Figure S3). These results confirm the presence of two accessory [4Fe-4S] clusters in the F-domain of ToHydA, as expected for M2-type hydrogenases. With methyl viologen (MV) as an electron mediator, To-HydA WT exhibits *in-vitro* H_2_ production activity of approximately 100 μmolH_2_.min^-1^.mg^-1^ (SI Figure S4A) which is comparable with CpIII.^21^ As the *T. oceani* is a thermophilic bacterium, *in-vitro* H_2_ production is measured at 60°C which results in approximately 2.5-3-fold increase in activity (SI Figure S4A). In addition, protein film electro-chemistry was used to evaluate the catalytic properties of ToHydA WT in both H^+^ reduction and H_2_ oxidation processes. Firstly, the cyclic voltammogram obtained for To-HydA WT shows thermodynamic reversibility with no detectable overpotential (SI Figure S5). Note that, the cyclic voltammogram of ToHydA WT also reveals inactivation at about -0.18 V (vs SHE at pH 7), tentatively indicating the O_2_ stability of ToHydA WT. Notably, this inactivation potential is about 100 mV higher than those observed in CbA5H and CpIII.^21,22,44^

### 2.2. The resting state of ToHydA WT is H_inact_ instead of H_ox_

In order to investigate various states of H-cluster in To-HydA WT, Attenuated Total Reflectance-Fourier Transform Infrared (ATR-FTIR) spectroscopy is employed in this study. ToHydA WT dried on the ATR crystal (referred to as the as-isolated state) exhibits several states that differ from those generally observed in Group A hydrogenases (Figure 2A). For FTIR spectroscopy, we used ToHydA WT protein containing 2 mM NaDT. However, as shown in SI Figure S6, the spectra of as-isolated ToHydA WT with and without 2 mM NaDT are highly comparable. The peak observed at 1826 cm^-1^ attributed to the bridging CO ligand, and the peaks at 1975 cm^-1^ and 1996 cm^-1^ corresponding to the terminal CO ligands, in the FTIR spectra of ToHydA WT, closely resemble the previously reported H_ox+1_ state of CpIII by Artz et al.^21^ Upon O_2_ treatment, the ToHydA WT predominantly accumulates the three CO ligand peaks mentioned earlier, the resulting state is referred to as the H_inact_ state (Figure 2A). Notably, no observable H-cluster degradation occurs during the O_2_ purging process, as also reported for CbA5H.^44,47^ The FTIR bands for the H_inact_ state in ToHydA WT are slightly shifted to lower wavenumbers (red-shifted) compared to CbA5H, although the position of the peaks clearly indicates the [Fe_p_(II)-Fe_d_(II)]_H_ electronic configuration of the H-cluster, likewise the H_inact_ state of CbA5H.^52^ The slight red shift in wavenumbers may be attributed to the different protein scaffolds surrounding the H-cluster in CbA5H and ToHydA. Surprisingly, enrichment of H_inact_ was also achieved upon N_2_ treatment (Figure 2A) that differs from Group A hydrogenases. The unusual formation of H_inact_ under N_2_ purging indicates that the resting state of ToHydA is the H_inact_ state and interestingly, not the H_ox_ state as reported for Group A hydrogenases.^7,53^ Leaving aside the predominant H_inact_ state, the as-isolated state of ToHydA WT exhibits a few peaks between 1884 cm^-1^ to 1947 cm^-1^ that might be associated with the H_ox_ state and the reduced states.

**Figure 2.**
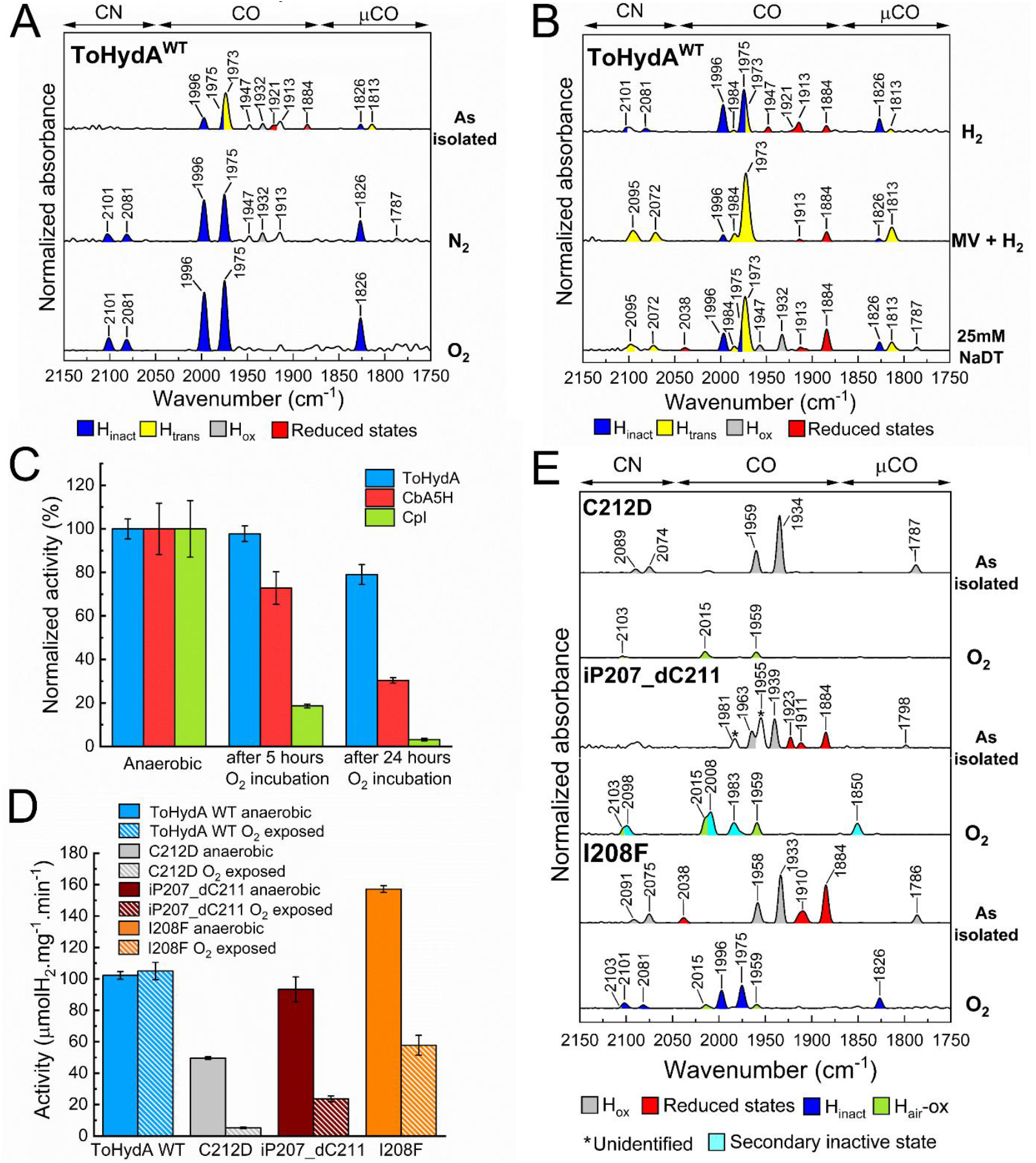
**A**. FTIR spectra of ToHydA WT in the as-isolated state (upper panel), and after N_2_ (N_2_ 8L/min, middle panel) and O_2_ (air 2L/min, lower panel) purging. B. FTIR spectra of ToHydA WT with H_2_ (5 L/min) exposure (upper panel), MV and H_2_ (H_2_ 5 L/min, protein sample without NaDT, middle panel), and with 25 mM NaDT. All gas purging experiments started from as-isolated state. The spectra of the proteins are normalized to the second amide band (1535-1545 cm^−1^). 0.4 - 0.5 mM protein sample was prepared in 100 mM Tris/HCl buffer (pH 8.0) containing 2 mM NaDT (otherwise mentioned). C. H_2_ production activities of ToHydA, CbA5H and CpI anaerobic and after 5h and 24h O_2_ incubation (replicates are shown in SI Figure S8). For O_2_ incubation, 10 μL of 15-20 mg/mL protein samples in 100 mM Tris-HCl buffer (pH8) containing 2mM NaDT were stored under air for 5 and 24 hours at 8°C. H_2_ production was measured at 37°C. D. *In-vitro* H_2_ production activity of ToHydA WT and variants before and after 20 minutes O_2_ incubation (replicates are shown in SI Figure S11). For O_2_ incubation, 6 μL of 25-30 mg/mL protein samples in 100 mM Tris-HCl buffer (pH8) containing 2mM NaDT were exposed to air on ice (at 4°C) for 20 minutes. H_2_ production was measured at 37°C. E. FTIR spectra of ToHydA variants in the as-isolated state and after O_2_ exposure. All gas purging experiments started from as-isolated state. The spectra of the proteins are normalized to the second amide band (1535-1545 cm^−1^). 0.4 - 0.5 mM protein sample was prepared in 100 mM Tris/HCl buffer (pH 8.0) containing 2 mM NaDT.

ToHydA was then treated with H_2_-saturated aerosol; however, the spectrum remains largely unchanged, suggesting ToHydA barely reacts to H_2_ (Figure 2B). To accelerate the response of ToHydA WT to H_2_, MV was used as an electron acceptor. For this experiment, first NaDT was removed from the protein sample to ensure that the observed reduction was due to H_2_ and not NaDT. The resulting FTIR spectra show an H-cluster state with a peak at 1813 cm−^1^ corresponding to the bridging CO ligands and a very intense band at 1973 cm−^1^ (Figure 2B). The consistent changes of peaks at 1973 cm−^1^ and 1813 cm−^1^ allowed us to assign them into a different state, together with the peaks of 2095 cm−^1^ and 2072 cm−^1^ for CN^-^ ligands. In addition, the peak of 1984 cm−^1^ could be attentively attributed to the second terminal CO of this new state. Given the slight red-shifted pattern (2-13 cm^-1^) of this state in comparison to H_inact_, we assign this state as H_trans_ state with the configuration of [4Fe]H^+^- [Fe_p_(II)-Fe_d_(II)]_H_ in which the [4Fe]_H_ is reduced comparing the H_inact_ state. The hydride-bound state, H_hyd_, shares similar peak positions in the FTIR spectrum and has the same electronic state as H_trans_.^25,54^ To rule out the possibility of the H_hyd_ state in this case, we applied the primary isotope effect by exchanging the buffer from H_2_O to D_2_O followed by D_2_ purging, as the H_hyd_ state involves a terminal hydride. This H/D effect was observed as a shift in the peak position of the bridging CO in the FTIR spectrum as demonstrated before for the H_hyd_ state.^54–57^ However, no noticeable changes of the IR frequence of the bridging CO ligand was observed, confirming that the state is H_trans_ rather than H_hyd_ (SI Figure S7). Formation of the H_trans_ state in ToHydA can also be induced with 25 mM sodium dithionite (NaDT), alongside the H_inact_, H_ox_, and reduced states (Figure 2B). In the as-isolated state, the distal CO bands of both H_inact_ and H_trans_ overlap, leading to a more intense band at 1975 cm^-1^ compared to the individual states (Figure 2A). The coexistence of these two states in the as-isolated ToHydA WT is evidenced by the appearance of two distinct bridging CO bands at 1826 cm^-1^ and 1813 cm^-1^.

### 2.3. ToHydA WT remains stable upon prolonged O_2_ exposure due to the presence of C212

To assess the stability of ToHydA under prolonged O_2_ exposure, which is highly relevant for sustainable energy conversion technologies, we measured the enzymatic activity of ToHydA WT using an MV-mediated *in-vitro* H_2_ production assay after 5 hours and 24 hours of O_2_ incubation (at 8°C) (Figure 2C; replicates are shown in SI Figure S8). For comparison, two Group A [FeFe]-hydrogenases CbA5H and CpI were used as controls. The results showed that at the end of 5 hours of O_2_ exposure, CbA5H and CpI retained approximately 75% and 18% of their H_2_ production activity, respectively. After 24 hours, their activity further decreased to approximately 30% and 3%, respectively (Figure 2C). In contrast, ToHydA retained nearly 100% of its H_2_ production activity after 5 hours of O_2_ exposure, with only a 20% reduction in activity at end of 24 hours of O_2_ exposure (Figure 2C). Additionally, The O_2_ stability of ToHydA WT remained unaffected by O_2_ incubation at higher temperatures (37°C and 60°C), as evidenced by its activity at both 37°C and 60°C (SI Figure S4B and S4C). This demonstrates the remarkable O_2_ stability of ToHydA under prolonged exposure. The intact [4Fe-4S]-clusters of ToHydA WT before and after O_2_ exposure were confirmed by UV-Vis spectroscopy (SI Figure S9A, S9B and S9C).

Prior to this study, the formation of the H_inact_ state was observed in three hydrogenases, in CbA5H through the coordination of intrinsic thiol of a cysteine while in DdH, and CrHydA1 through the extrinsic sulfide.^31,44^ In our infrared spectroscopic characterization, no extrinsic sulfide was used, therefore we hypothesize ToHydA enters the O_2_-protected H_inact_ state via a cysteine (presumably C212)-dependent binding to Fe_d_, as observed in CbA5H.^44^ To confirm this, a ToHydA C212D variant was produced (mutation site shown in SI Figure S10; primers used for QC-PCR are listed in Table 1; SDS-PAGE shown in SI Figure S2). Upon O_2_ exposure, the ToHydA C212D variant displayed severely reduced H_2_ production activity (Figure 2D; replicates are shown in SI Figure S11). The H_2_ production activity is further reduced under O_2_ exposure at higher temperatures (SI Figure S4B and S4C). The integrity of the [4Fe-4S] clusters in anaerobic and O_2_ exposed ToHydA C212D variant was confirmed through UV-Vis spectroscopy (SI Figure S9A, S9B and S9D). As we anticipated, in FTIR spectra, the signature peaks representing the H-cluster were mostly diminished in the presence of O_2_, and instead, three small bands appeared at higher wavenumbers (Figure 2E), closely resembling the degraded mononuclear H_air_-OX state as identified in Group D hydrogenases.^18^

### 2.4. The additional Cysteine is essential for the high O_2_-stability of ToHydA

In Group B (M2a) hydrogenases, the presence of an additional cysteine alongside the two functionally crucial and conserved cysteine residues near the H-cluster is particularly intriguing. We created a cysteine deletion variant of ToHydA by deleting the additional cysteine, abbreviated as ToHydA dC211 (mutation site shown in SI Figure S10; primers used for QC-PCR are listed in Table 1; SDS-PAGE shown in SI Figure S2). Although the dC211 ToHydA locally mimics the TSCCP motif of Group A hydrogenases, the deletion of C211 shortens the protein sequence by one residue, potentially causing significant mutational artifacts. This is reflected in a 70% reduction in the *in-vitro* H_2_ production activity compared to ToHydA WT (see SI Figure S12A). To circumvent these mutational artifacts, we revisited the sequence alignments of Group A and Group B (M2a) hydrogenases (see Figure 1A for the sequence alignments) to identify other differences that might counterbalance the effects of the C211 deletion. The sequence alignment reveals that Group A hydrogenases, such as CbA5H, CpI, and CrHydA1, contain a highly conserved proline residue insertion (P294 in CpI, P362 in CbA5H, and P164 in CrHydA1) in the upstream part of the TSCCP motif, comparing with Group B (M2a) hydrogenases, including ToHydA and CpIII. A more detailed analysis on 72 Group B (M2a) hydrogenases and 72 Group A hydrogenases obtained from HydDB^42^ supports this observation (SI Figure S1). This difference suggests that the presence of proline in the upstream part of the TSCCP motif in Group A hydrogenases may counter-balance the absence of the additional cysteine next to the proton-transporting cysteine. To explore this hypothesis, we designed a double variant of ToHydA by deleting the additional cysteine (C211) and simultaneously inserting a proline residue between F206 and M207, referred to as iP207_dC211 ToHydA (mutation sites shown in SI Figure S10; primers used for QC-PCR are listed in Table 1; SDS-PAGE shown in SI Figure S2). This variant is intended to closely resemble the proton-transporting cysteine bearing loop of Group A hydrogenases. The iP207_dC211 variant exhibits nearly 100% anaerobic H_2_ production activity (Figure 2D; replicates are shown in SI Figure S11), compared to To-HydA WT.

After O_2_ exposure, the *in-vitro* H_2_ production activity of ToHydA iP207_dC211 is reduced to approximately 30% of its anaerobic activity (Figure 2D; replicates are shown in SI Figure S11). The H_2_ production activity decreases to 5% of its anaerobic activity under O_2_ exposure at higher temperatures (SI Figure S4B and S4C). The integrity of the [4Fe-4S] clusters in anaerobic and O_2_ exposed ToHydA iP207_dC211 was confirmed by UV-Vis spectroscopy (SI Figure S9A, S9B and S9E). The FTIR spectrum of ToHydA iP207_dC211 shows several states upon O_2_ purging, with overall low peak intensity due to H-cluster degradation (Figure 2E). Notably, the H_inact_ state is not observed in the FTIR spectra under O_2_ conditions. Instead, peaks at 2015 cm^-1^ and 1959 cm^-1^, attributed to the H_air_-OX state, appear similar to the O_2_-treated ToHydA C212D variant. Compellingly, in the O_2_-treated To-HydA iP207_dC211 FTIR spectra, a peak at 1850 cm^-1^ corresponding to a bridging CO ligand, along with two peaks at 1983 cm^-1^ and 2008 cm^-1^ representing terminal CO ligands, appear that primarily indicates a partly intact H-cluster. This H-cluster state closely resembles with the previously reported O_2_-bound H_ox_-O_2_ state observed in the O_2_-treated C169A CrHydA1 variant (a proton transport-impaired variant),^58^ as well as the aqua or hydroxide bound state found in the N_2_-treated Group D TamHydS variants targeting the proton transfer pathway.^59^ After conducting additional experiments, we hypothesize that an aqua or hydroxide ligand coordination at the Fe_d_ can be the molecular origin of this non-canonical inactivation state similar to TamHydS variants (for details see SI Figure S13). Further studies are needed to clarify the nature of the observed inactivation state, including the putative role of water molecules. In the present study, this state is referred to as the secondary inactive state. The formation of this speculated inactive state might protect the H-cluster, as evidenced by the residual 30% H_2_ production activity in the *in-vitro* assay of O_2_-treated iP207_dC211 ToHydA. The ToHydA dC211 variant exhibits behavior similar to the iP207_dC211 variant under O_2_ treatment, as evidenced by a decrease in catalytic activity after O_2_ exposure (SI Figure S12A). Additionally, FTIR spectroscopy shows the accumulation of the H_air_-OX state and the secondary inactive state upon O_2_ purging (SI Figure S12B).

### 2.5. I208 in ToHydA controls O_2_ stability

Although the O_2_-stability of CbA5H and ToHydA both depends on the presence of the conserved cysteine residue (C367 in CbA5H and C212 in ToHydA), the molecular determinants modulating the formation of H_inact_ differ between these two enzymes. Previous studies have shown that in CbA5H, the LPA motif, comprising L363, P386, and A561 residues, plays a crucial role in forming H_inact_.^44^ Sequence alignment using Clustal Omega shows that the LPA motif of CbA5H is absent in ToHydA (SI Figure S14). Notably, the leucine (L363) residue in CbA5H is replaced by structurally similar isoleucine (I208) in ToHydA, suggesting that I208 could play a role in the O_2_-protection mechanism of ToHydA. To probe the role of I208 in ToHydA’s O_2_-stability, we replaced I208 with phenylalanine, a residue found in O_2_-sensitive CpI and CrHydA1 hydrogenases. Interestingly, the ToHydA I208F variant (mutation site shown in SI Figure S10; primers used for QC-PCR are listed in Table 1; SDS-PAGE shown in SI Figure S2) exhibits higher H_2_ production activity comparing WT enzyme under anaerobic conditions. However, upon exposure to O_2_, the H_2_ production activity of I208F significantly decreased to approximately 35% of its anaerobic activity (Figure 2D; replicates are shown in SI Figure S11). Interestingly, unlike iP207_dC211 the O_2_-induced degradation in the I208F variant was found to be temperature-independent as evidenced by the consistent 35% H_2_ production activity of anaerobic protein after O_2_ exposure at 37°C and 60°C (SI Figure S4B and S4C). The presence of intact [4Fe-4S] clusters in I208F variant before and after O_2_ exposure was confirmed by UV-Vis spectroscopy (SI Figure S9A, S9B and S9F). Under constant O_2_ flow, the as-isolated I208F variant of ToHydA showed a significantly reduced intensity of the H_inact_ state compared to the WT (Figure 2E). The decreased intensity of H_inact_ peaks suggests some degradation of the H-cluster, with a smaller portion forming H_air_-OX, similar to the O_2_-exposed ToHydA C212D variant. These outcomes suggest that formation of H_inact_ is partially hindered in I208F and emphasize the role of the I208 residue in ToHydA for O_2_-stability and H_inact_ state formation. To gain a more detailed understanding, we conducted extensive MD simulations of the ToHydA WT along with iP207_dC211 and I208F variants, as presented below.

### 2.6. The hydrophobic cluster around I208

The conformational flexibility of the C212-bearing loop in ToHydA is fundamental for the intrinsic thiol mediated inactivation mechanism that protects against O_2_ exposure. To understand the loop dynamics at an atomic level, MD simulations were conducted using the AlphaFold-predicted models of WT ToHydA and its variants (SI Figure S15 and the equilibrated structure of WT ToHydA is provided as supporting information). A total of 1500 ns of MD trajectories were analyzed for each system (see SI detailed MD methods). A key parameter in this protection mechanism is the distance between the sulfur atom of C212 (C212:S_γ_) and the Fe_d_ atom in the [2Fe]_H_ cluster, as this distance determines how easily C212 can reach Fe_d_ in an O_2_-rich environment. Figure 3A shows the distribution of this distance in the MD simulations. In the WT enzyme, the distribution indicates that C212 remains close to Fe_d_, while in the I208F and iP207_dC211 variants, C212 is located further away from Fe_d_. The distributions also provide insights into the dynamics of the C212 residue. The WT enzyme exhibits a broad distance distribution, ranging from 4 Åto 7 Å, suggesting significant flexibility in the C212-bearing loop. In contrast, the iP207_dC211 variant shows a narrower distribution (5.5 Åto 7 Å), indicating reduced flexibility. The I208F variant has a broader distribution (4 Åto 8 Å), suggesting that C212 remains more dynamic than the WT. A correlation between the C212:S_γ_…Fe_d_ distance distribution and the root mean square fluctuation (RMSF), a measure of protein flexibility, was also observed. The RMSF of C212 is similar in both the WT and I208F variants, while it is lowest in the iP207_dC211 variant (Figure 3B). This reduced flexibility in the iP207_dC211 variant aligns with the narrower distance distribution and results in a nearly immobile C212, which impedes the H_inact_ formation in the presence of O_2_. Interestingly, the RMSF of F208 in the I208F variant is higher than that of I208 in the WT.

**Figure 3.**
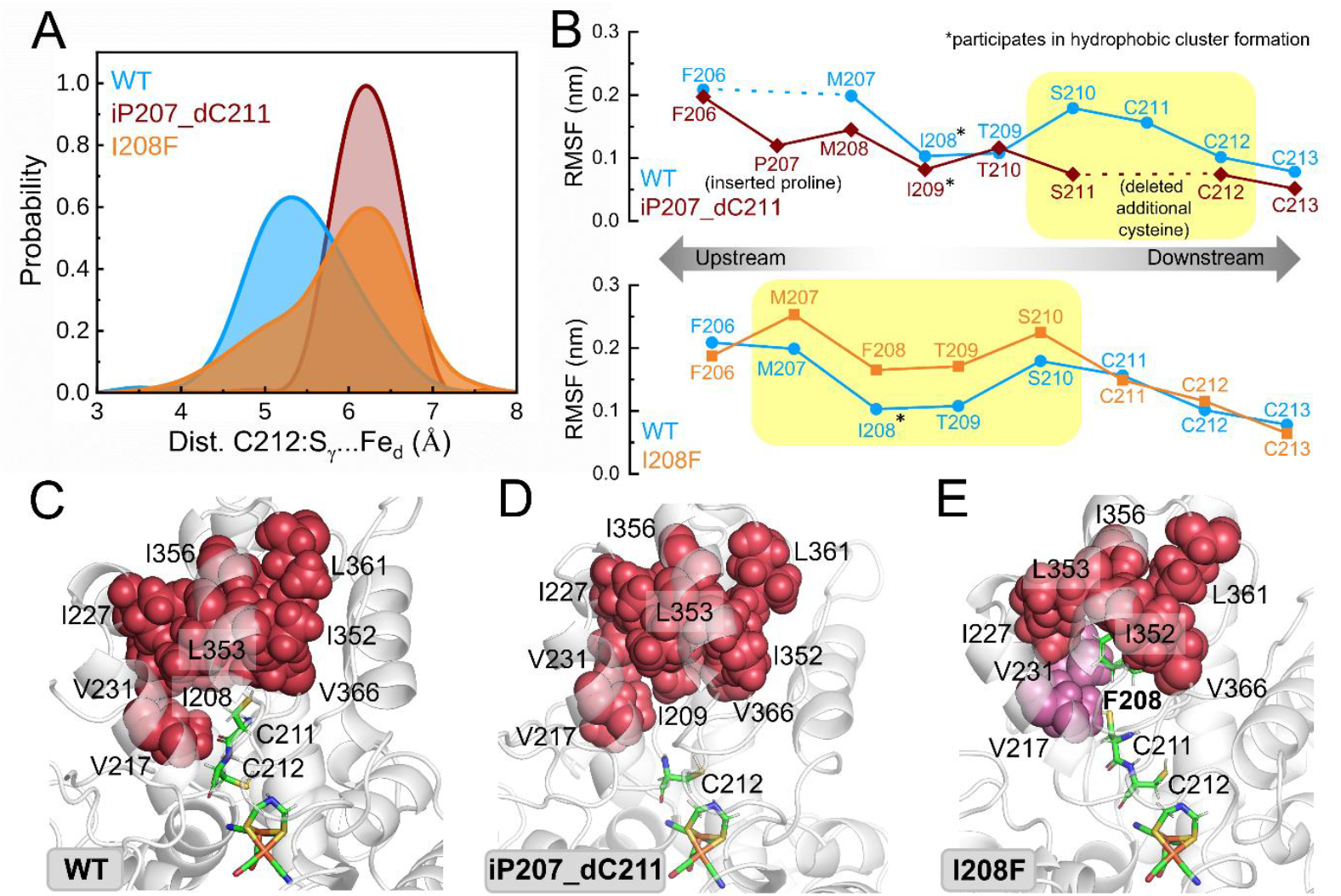
MD simulations of ToHydA WT and its variants. A. Distance distribution between C212:S_γ_ and Fe_d_ in ToHydA WT and variants. B. The root means square fluctuation (RMSF), a measure of protein structural flexibility, is shown for the C212 bearing loop. The differences in RMSF are highlighted by the yellow box. C-E. The hydrophobic cluster in ToHydA WT, centered around the C212-bearing loop, involving Val (V), Ile (I), and Leu (L) residues is illustrated by red van der Waals spheres. This hydrophobic cluster is preserved in the iP207_dC211 variant. However, in the I208F variant, this cluster splits into two separate clusters, depicted in red and magenta.

To gain further insights, our aim was to pinpoint distinct structural elements influencing protein folding and stability. Using the ProteinTool webserver,^60^ we identified a previously unrecognized hydrophobic cluster around residue I208 in WT ToHydA (Figure 3C). This cluster includes I208, V217, I227, V231, I352, L353, I356, L361, and V366, the positions mostly accommodated by hydrophobic residues such as Ile, Leu, Val, or Met across Group B (M2a) [FeFe]-hydrogenases. This hydrophobic cluster contributes to structural rigidity, as is indicated by the low RMSF of I208 in WT ToHydA (Figure 3B), which might help position the C212-bearing loop, thereby bringing the C212 side chain into close proximity to the Fe_d_ center under O_2_-rich conditions. A similar hydrophobic cluster is also present in the O_2_-stable CbA5H (centered around L364), suggesting that it could be an important feature in the H_inact_ forming hydrogenases (SI Figure S16A). In the I208F variant, replacing isoleucine with phenylalanine at position 208 disrupts this hydrophobic cluster by splitting it into two smaller clusters (Figure 3E), which leads to enhanced fluctuations at position 208 (Figure 3B, lower panel). This splitting prevents the C212-bearing loop from attaining an optimal position, leading to increased C212:S_γ_…Fe_d_ distances (Figure 3A) and reduced efficiency of H_inact_ formation in the I208F variant. Similar disruptions by phenylalanine residues are also noted in the O_2_-sensitive Group A CrHydA1 and CpI hydrogenases (SI Figure S16B and S16C). In the iP207_dC211 variant, I209 forms a hydrophobic cluster with neighboring isoleucine, leucine, and valine residues (V217, I277, V231, I352, L353, I356, L361, and V366), similar to WT ToHydA (Figures 3C and 3D). This cluster holds the C212-bearing loop at position 209, as evidenced by the low RMSF of I209 in the iP207_dC211 variant (or of the corresponding I208 in the WT). The insertion of proline residue also reduces the protein’s flexibility in the upstream part of the C212-bearing loop. Meanwhile, the shortening of the loop due to the deletion of cysteine (C211 in ToHydA WT) decreases overall protein flexibility. Further inspection of the simulations revealed that the backbone of the C212-bearing loop in WT and variants forms a dynamic hydrogen bond network with the backbones of two neighboring loops (SI Figure S17). The WT and I208F variant form up to three hydrogen bonds with neighboring loops, while this number increases to five in the iP207_dC211 variant, stiffening this part of the protein and restricting the mobility of the C212-bearing loop in the iP207_dC211 variant compared to the WT.

## Discussion

This study investigates the O_2_ stability of Group B (M2a) ToHydA hydrogenase through a combination of biochemical analyses, FTIR spectroscopy, and atomistic MD simulations. FTIR spectroscopy together with site-directed mutagenesis experiments confirms the formation of the H_inact_ state through involvement of C212 in the O_2_-rich environment, similar to that seen in Group A CbA5H hydrogenase.^44^ ToHydA has significantly improved O_2_ stability compared to CbA5H, even after prolonged exposure. It is particularly noteworthy that ToHydA fully recovers its *in-vitro* H_2_ production activity after O_2_ exposure, matching the activity observed under anaerobic conditions. Based on data from CbA5H, we proposed a model for O_2_-protection in ToHydA which aligns with our observations in ToHydA as shown in SI Figure S18.

Unlike Group A hydrogenases, which typically adopt the H_ox_ resting state characterized by the oxidation states of [4Fe]H^2+^-[Fe_p_(II)-Fe_d_(I)]_H_ with an open coordination site at Fe_d_ which is essential for the initiation of the catalytic cycle,^53^ ToHydA WT exhibits an unusual resting state, H_inact_. In other [FeFe]-hydrogenases, this state is known to have the electronic configurations of [4Fe]H^2+^-[Fe_p_(II)-Fe_d_(II)]_H_, with the Fe_d_ site being coordinated by a thiol ligand.^21,31,44^ Since the flexibility of C212, along with the C212:S_γ_…Fe_d_ distance, is crucial for efficient H_inact_ formation, this dynamic behavior directly correlates with the extent of H_inact_ formation. The propensity for H_inact_ formation in ToHydA WT, even under N_2_ exposure, suggests a prolonged proximity of the C212:S_γ_ to the Fe_d_ site, as evidenced by the C212:S_γ_…Fe_d_ distance distribution observed in MD simulations of ToHydA WT. This further stabilizes the H_trans_ state which is obtained from the H_inact_ state upon reduction of the [4Fe]_H_ cluster by addition of a reducing agent such as NaDT. Although the H_trans_ state was previously identified in Group A DdH hydrogenase as an intermediate during H_inact_ formation, where an extrinsic sulfide ligand binds to the Fe_d_ site,^15,31,32^ this state is not observed in Group A CbA5H hydrogenase, likely due to its transient nature.

Despite the H_ox_ state bands being rarely detected in the FTIR spectra of ToHydA WT, the enzyme continues to exhibit *in-vitro* H_2_ production activity. This suggests that the H_ox_ state transiently forms prior to initiating the catalytic cycle which can be achieved through an electronic rearrangement of the H_trans_ state followed by the relocation of the C212 thiol group. Notably, the overall oxidation state of the H-cluster, encompassing both [2Fe]_H_ and [4Fe]_H_, remains unchanged during the transition between the H_trans_ and H_ox_ states (Figure 4A), which may imply a spontaneous transition from the H_trans_ state to the H_ox_ state during catalysis. This hypothesis of a transient transition from H_trans_ to H_ox_ is directly supported by the complete or partial accumulation of the H_ox_ state in the as-isolated C212D, I208F, and iP207_dC211 variants (Figure 2E), where the formation of the H_inact_ state via involvement of C212 is either impaired or severely hindered. It is evident that ToHydA favors the [Fe_p_(II)-Fe_d_(II)]_H_ electronic configuration over [Fe_p_(II)-Fe_d_(I)]_H_, although the underlying reason for this preference remains unidentified. However, this inclination toward the [Fe_p_(II)-Fe_d_(II)]_H_ configuration may enhance the O_2_ stability of ToHydA significantly, even after prolonged exposure to O_2_, in comparison to CbA5H.

**Figure 4.**
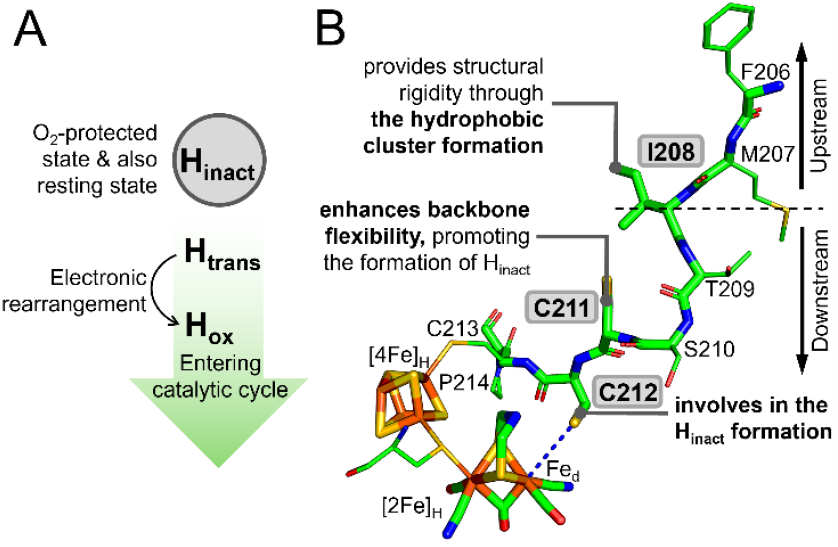
A schematic representation of A. the different states of the H-cluster of ToHydA and, B. the roles of key residues within the TSCCCP motif and hydrophobic cluster of ToHydA as observed in this study.

Sequence alignment of Group B (M2a) hydrogenases, including ToHydA, reveals a marked difference from Group A hydrogenases, notably due to the presence of an additional cysteine residue adjacent to a pair of conserved, catalytically essential cysteine residues near the H-cluster. The To-HydA dC211 variant (C211 deletion variant) exhibits significantly reduced catalytic activity which further highlights the lack of a proline residue upstream of the TSCCCP motif (Figure 1A). Remarkably, the catalytic activity of the iP207_dC211 variant (in which proline residue was introduced at position 207 to compensate for the deletion of C211), as measured by H_2_ production, is restored to WT levels. This finding strongly suggests that the additional C211 cysteine residue is not directly involved in catalysis. However, unlike the ToHydA WT, the iP207_dC211 variant fails to protect the H-cluster from O_2_-induced degradation via the canonical H_inact_ formation pathway as evident from the sharply reduced activity of the iP207_dC211 variant following O_2_ exposure, compared to WT. FTIR spectra of the O_2_-exposed iP207_dC211 suggests that the H-cluster appears to be partially protected. As shown in SI Figure S19, water molecules near the [2Fe]_H_ are likely to block the open coordination site of the H-cluster. This suggests that water molecules might serve as an alternative mechanism of protecting the H-cluster; though the hypothesis requires further investigation. This alternative protection mechanism has not been previously observed in catalytically active Group A hydrogenases, where O_2_ exposure does not induce cysteine-mediated H_inact_ formation. However, a similar state called O_2_-bound state has been reported in catalytically inactive Group A variants with disrupted proton transport pathways.^58^ Moreover, analogous inactive states are also detected in the ToHydA dC211 and C212A variants. The previous study by Fasano et al.^22^ on Group B (M2a) CpIII hydrogenase also reported a second inactive state (distinct from the H_inact_ state) in a cysteine-deleted CpIII variant (analogous to the ToHydA dC211 variant). Notably, the secondary inactive state does not occur in WT ToHydA, likely due to the close proximity of a proton-transporting cysteine residue to the Fe_d_, which prevents water molecule coordination and favors the formation of the canonical H_inact_ state.

MD simulations demonstrate that the isoleucine residue of the C212-bearing loop in the WT forms a rigid hydrophobic cluster comprised of valine, isoleucine, and leucine residues. The formation of hydrophobic clusters has often been identified to influence the protein folding and stability, which is also associated with reduced flexibility in specific regions of folded proteins.^61,62^ The hydrophobic clustering in ToHydA WT immobilizes I208 and restricts the overall mobility of the loop at this position, effectively dividing the C212-bearing loop into two distinct segments (Figure 4B). The downstream portion of I208, containing the TSCCCP sequence in WT, retains high flexibility, which enhances the C212 mediated Fe_d_ site protecting efficiency and promotes the formation of the H_inact_ state. However, in the case of iP207_dC211 variant, the deletion of a cysteine from this segment reduces its flexibility, increasing the C212 distance from the Fe_d_ site, which significantly impairs H_inact_ formation. Furthermore, the proline insertion in the upstream segment of the iP207_dC211 variant (intended to compensate for the cysteine deletion in the downstream segment) does not improve the C212-mediated H_inact_ formation, owing to the anchoring of the isoleucine residue (I209 in case of iP207_dC211) through hydrophobic cluster formation. Despite the peptide length of the C212-bearing loop remaining identical to the WT, H_inact_ formation in the iP207_dC211 variant is hindered. Interestingly, similar hydrophobic cluster formations are observed in O_2_-stable Group A CbA5H hydrogenase but not in O_2_-sensitive Group A hydrogenases like CpI or HydA1, where a phenylalanine residue disrupts the cluster. This phenylalanine-mediated disruption is also seen in the ToHydA I208F variant, leading to a displacement of the C212-bearing loop from its optimal position, as indicated by the increased C212 distance from the Fe_d_ site and elevated loop flexibility at position 208. Consequently, this disruption hinders efficient H_inact_ formation, leading to H-cluster degradation upon O_2_ exposure. Altogether, in To-HydA the rigid hydrophobic cluster and flexible TSCCCP motif collectively assists the O_2_-stability mechanism. However, these characteristics may not necessarily represent an evolutionary adaptation to oxygen exposure as the host organism is strictly anaerobic.

## Conclusions

In conclusion, this study elucidates several critical factors contributing to the O_2_-stability of the Group B (M2a) [FeFe]-hydrogenases ToHydA through biochemical assays, ATR-FTIR spectroscopy, site-directed mutagenesis and MD simulation. Our findings reveal that ToHydA forms H_inact_ state that protects the H-cluster from O_2_-induced degradation and demonstrates almost 80% *in-vitro* H_2_-production activity, after 24 hours O_2_ exposure. ToHydA also forms H_inact_ state as its auto-oxidized resting state. The intrinsic sulfide-dependent H_trans_ state during the transition from H_inact_ to active states is observed in ToHydA, notably independent of extrinsic sulfide. Furthermore, sequence alignments of Group A and Group B (M2a) [FeFe]-hydrogenases reveal the presence of an additional cysteine residue adjacent to the proton-transporting cysteine and a missing proline residue upstream of the two former cysteines. The additional cysteine residue in ToHydA is not directly involved in catalytic activity, but it is essential for O_2_-stability which increases the flexibility of the TSCCCP motif. Atomistic MD simulations identify a unique hydrophobic cluster centered around the proton-transporting C212-bearing loop in To-HydA. The position I208, at the center of the hydrophobic cluster, influences the loop dynamics and is crucial for the O_2_-stability of ToHydA. The rigid hydrophobic cluster, in combination with the flexible TSCCCP motif, enhances the O_2_-stability mechanism in ToHydA. Altogether, these findings not only advance the understanding of O_2_ stability mechanisms in [FeFe]-hydrogenases but also pave the way for the development of more efficient and durable biocatalysts for sustainable energy conversion technologies.

## ASSOCIATED CONTENT

### Supporting Information

The supporting information is available free of charge at http://pubs.acs.org. The supporting information includes experimental and simulation details, sequence analysis, SDS-PAGE of ToHydA WT and variants, ICP-OES results, UV-Vis spectroscopic data, H_2_-production at high-temperatures, electrochemistry data, H_2_ production activity, isotope-exchange FTIR spectra, H_2_ production activity and FTIR spectra of dC211, additional experiments regarding secondary inactive state, several biochemical assay replicate information, and additional MD analysis. The Clustal Omega sequence alignment of 144 [FeFe]-hydrogenases is uploaded as a separate pdf file. The equilibrated structure of ToHydA WT is provided as a separate pdb file.

## AUTHOR INFORMATION

**Notes**

The authors declare no competing financial interest.

## ACKNOWLEDGMENT

S.G. thanks Deutscher Akademischer Austauschdienst (DAAD) for funding her doctoral scholarship. This project received funding from the Deutsche Forschungsgemeinschaft (DFG) under Germany’s Excellence Strategy – EXC 2033 – 390677874 – RESOLV. J.D. thanks the funding from DFG (project number 461338801). We are grateful to Shanika Yadav and Ulf-Peter Apfel (Faculty for Chemistry and Biochemistry, Ruhr University Bochum) for synthesizing and providing the [2Fe]_MIM_ for *in-vitro* maturation.

