## Supporting Information for "Protein dynamics affect O_2_-stability of Group B [FeFe]-hydrogenase from *Thermosediminibacter oceani*"

### Table of Contents

Experimental details ... 3
Sequence ... 3
Site-directed mutagenesis ... 3
Expression and purification ... 3
*In-vitro* maturation ... 3
Iron quantification by inductively coupled plasma-optical emission spectroscopy (ICP-OES) ... 4
UV-visible (UV/Vis) spectroscopy ... 4
$H_2$ production assay ... 4
Protein film electrochemistry ... 5
Attenuated total reflectance-Fourier transform infrared (ATR–FTIR) spectroscopy ... 5
Molecular dynamics simulation details ... 5
Supporting figures and tables ... 7
Figure S1. Comparison of proton transporting cysteine bearing loop of group A and group B hydrogenases ... 7
Figure S2. SDS PAGE of ToHydA WT and variants after purification ... 8
Figure S3. Fe content of ToHydA WT ... 8
Figure S4. *In-vitro* $H_2$ production activity of ToHydA WT and variants at elevated temperatures ... 9
Figure S5. Electrochemistry ... 9
Figure S6. FTIR spectra of as-isolated ToHydA WT with 2mM NaDT and without NaDT ... 10
Figure S7. The "$H_2$–$N_2$" ("$D_2$-$N_2$" in case of $D_2O$) difference FTIR spectrum of ToHydA in $H_2O$ and $D_2O$ ... 10
Figure S8. *In-vitro* $H_2$ production activity of ToHydA$^{WT}$, CbA5H$^{WT}$, and CpI$^{WT}$ ... 11
Figure S9: UV-Vis spectra of ToHydA WT and its variants. ... 12
Figure S10. Sequence of the C212-bearing loop for ToHydA WT and variants ... 13
Figure S11. *In-vitro* $H_2$ production activity of ToHydA WT and variants ... 14
Figure S12. Biochemical and ATR-FTIR spectroscopic characterization of ToHydA dC211 ... 15
Figure S13. ATR-FTIR spectra of variants ToHydA C212A and ToHydA iP207_dC211 ... 16
Figure S14. Sequence Comparison of LPA motif of CbA5H with ToHydA and CpI ... 16
Figure S15. ToHydA WT whole protein representation ... 17
Figure S16. Hydrophobic cluster formation in CbA5H, CrHydA1 and CpI ... 17
Figure S17. probability of hydrogen bond formation between the C212-bearing loop and neighboring loops ... 18
Figure S18: Proposed model for $H_{inact}$ formation in ToHydA ... 18
Figure S19. Water molecules near active site of ToHydA iP207_dC211 in MD simulation ... 19
Table S1. Generation of site-directed mutagenesis variants of ToHydA. Primers used for QuikChange PCR (QC-PCR) to generate variants of ToHydA. ... 19

References........................................................19

### Experimental details

#### Sequence

An *Escherichia coli* codon-optimized ToHydA sequence was designed using the sequence available in the manually curated hydrogenase database, HydDB[1] (WP_013276251.1) as a template. The ToHydA gene is ligated to the pET21b vector with NdeI and SalI restriction sites. The strep-tag II (peptide sequence of WSHPQFEK) is added at the N-terminal of the ToHydA sequence with a AS linker between the sequence and the tag.

#### Site-directed mutagenesis

Hydrogenase genes with different codons were generated employing QuikChange PCR, following the previously described protocol.[2] Expression plasmids were amplified with mismatch primers (SI Table S1). The resulting products were treated with DpnI endonuclease before being transformed into *E. coli* strain DH5α using the heat shock method. All DNA constructs were subsequently confirmed through sequencing.

#### Expression and purification

The *E. coli* strain BL21 (DE3) ΔiscR[3] was transformed with the pET21b expression plasmid harboring a codon-optimized ToHydA gene to express the apo form of ToHydA, which lacks the $[2Fe]_H$ subcluster.[4] The expression and purification processes were conducted in the absence of the hydrogenase maturases HydE, HydF, and HydG, under strictly anaerobic conditions as previously described.[4–6] The protein was purified by affinity chromatography using Strep-Tactin high-capacity resin (IBA GmBH). Protein concentration was measured by the Bradford assay,[7] and purity was confirmed by SDS-PAGE (SI Figure S2).[8] The purified proteins were stored at −80 °C in 100 mM Tris-HCl buffer (pH 8) containing 2 mM sodium dithionite (NaDT).

#### *In-vitro* maturation

To reconstitute the active protein, apoproteins were incubated on ice for 1 hour with a 10-fold molar excess of the artificially synthesized $[2Fe]_H$ cofactor mimic ($[Fe_2[\mu\text{-}(SCH_2)_2NH]\text{-}(CN)_2(CO)_4]^{2-}$)[9] in 100mM potassium phosphate ($K_2HPO_4/KH_2PO_4$) buffer (pH 6.8) with added 2mM sodium dithionite (NaDT), following the previously described method.[10] After incubation, holoproteins were separated from the excess $[2Fe]_H$ by size exclusion chromatography using a NAP 5 column (GE Healthcare) and the maturated holo-proteins were stored at −80 °C in 100 mM Tris-HCl buffer (pH 8) containing 2 mM NaDT.

Iron quantification by inductively coupled plasma-optical emission spectroscopy (ICP-OES)

Iron quantification was conducted using a Perkin–Elmer Optima 2100DV inductively coupled plasma optical emission spectrometer (ICP-OES) (Perkin–Elmer, Fremont, CA, USA). Protein samples underwent overnight wet digestion in a 1:1 solution of 65% nitric acid (Suprapur, Merck, Darmstadt, Germany) at 100 °C. Before ICP-OES analysis, the digested samples were diluted tenfold with water. Multielement standard solutions XII and XVI (Merck) were used as reference standards.

UV-visible (UV/Vis) spectroscopy

UV/Vis spectroscopy was carried out using an Shimadzu Spectrometer. Protein solutions at concentrations 10 µM were prepared in 100 mM Tris-HCl buffer (pH 8.0) without NaDT to ensure the enzyme remained in its oxidized state. For anaerobic sample preparation, 200 µM hexaammineruthenium(III) chloride (HAR) was added to oxidize all the [Fe-S] clusters. For aerobic sample preparation, anaerobic protein samples were exposed to air for 20 minutes to fully oxidize all [Fe-S] clusters.

$H_2$ production assay

To quantify *in-vitro* $H_2$ production activity, a reaction mixture was prepared in an 8 mL air-tight glass vessel, containing 10 mM methyl viologen (MV) as the electron mediator, 100 mM NaDT as the sacrificial electron donor, and 400 ng (for CpI and CbA5H)/800-1600 ng (ToHydA WT and variants) of holoenzyme in 100 mM potassium phosphate buffer ($K_2HPO_4$/$KH_2PO_4$) at pH 6.8. The mixture was degassed with 100% argon for 5 minutes and then incubated at 37°C (or 60°C for experiments in SI Figure S4A and S4C) for 30 minutes. The amount of $H_2$ produced in the headspace was measured using gas chromatography (Shimadzu).[11]

To examine the $O_2$ stability of ToHydA, CbA5H, and CpI, 10 µL of 15-20 mg/mL protein samples in 100 mM Tris-HCl buffer (pH8) containing 2mM NaDT were stored under air for 5 and 24 hours at 8°C. Protein dilution were prepared in anaerobic buffer and $H_2$ production was measured at 37°C by following the above-mentioned procedure (each time the proteins stored anaerobically measured as a control, Figure 2C).

To assess $H_2$ production of ToHydA WT and variants after $O_2$ incubation, 6 µL of 25-30 mg/mL protein samples in 100 mM Tris-HCl buffer (pH8) containing 2mM NaDT were exposed to air on ice (at 4°C) for 20 minutes (Figure 2D). At the end of the incubation period, the protein solutions were prepared in anaerobic buffer, promptly added to the reaction mixture to stop further $O_2$-induced damage. $H_2$ production was measured at 37°C following the standard *in-vitro* assay.

To measure $H_2$ production of ToHydA WT and variants after $O_2$ incubation at elevated temperatures (SI Figure S4B and S4C), 6 µL of 25-30 mg/mL protein samples in 100 mM Tris-HCl buffer (pH8) containing 2mM NaDT were exposed to air at elevated temperatures (37°C or 60°C), followed by $H_2$ production measured at their respective incubation temperatures (37°C or 60°C).

##### Protein film electrochemistry

Cyclic voltammetry experiments were performed under strictly anoxic conditions at 10°C in a sealed electrochemical cell. The cell was equipped with a hydrogen line for $H_2$ oxidation studies and operated using a PalmSens4 potentiostat with PStrace software. The working electrode was a rotating pyrolytic graphite edge disk at 3000 rpm, with a platinum wire counter-electrode and an Ag/AgCl (3M KCl) reference electrode. Potentials were corrected to the SHE scale ($E_{SHE} = E_{Ag/AgCl} + 0.217$ V at 10°C). Prior to measurements, 2–4 µL of holo [FeFe]-hydrogenase (~10 µM) was applied to the polished working electrode surface, incubated for 5 minutes, and rinsed. Electrochemical experiments were then conducted in 100 mM potassium phosphate buffer ($K_2HPO_4/KH_2PO_4$) buffer (pH 7) with 0.1 M NaCl.

##### Attenuated total reflectance-Fourier transform infrared (ATR−FTIR) spectroscopy

ATR-FTIR spectroscopy was conducted using a Bruker Tensor II spectrometer (Bruker Optik, Germany) equipped with a 9-reflection ZnSe/Si crystal (Microm ATR Vision, Czitek). All the measurements were performed under anaerobic conditions (1.5% $H_2$, 98.5% $N_2$) at 25°C. The FTIR spectra were recorded over a range of 4000 to 1000 $cm^{-1}$ with a resolution of 2 $cm^{-1}$. 4 µL of 0.4-0.5 mM protein sample stored in 100 mM Tris-HCl buffer (pH8) containing 2mM NaDT was spread onto the ATR crystal. All FTIR protein samples contained 2 mM NaDT unless stated otherwise.

The "as-isolated" ToHydA spectrum was prepared by drying under a tent atmosphere without an external gas stream. For gas-flushing experiments, the as-isolated sample was partially dried until the characteristic H-cluster bands were faintly visible, indicating a similar spectrum to the "as-isolated" spectrum but at lower intensity while maintaining hydration. To prevent further drying, the sample was then purged with a hydrated gas stream using $N_2$ (8 L/min), $O_2$ (2 L/min), or $H_2$ (5 L/min) to prevent further drying.

For the experiment in Figure 2B (middle panel), NaDT was removed from the matured sample. After drying the protein sample on the ATR crystal (procedure similar to as-isolated spectra), a 4 µL film of 0.2 mM MV was applied, followed by partial drying and purging with hydrated $H_2$ at 5 L/min. For chemical treatments such as NaDT and HAR, 4 µL of 25 NaDT or 4 µL of 100 mM HAR was applied to the dried protein film (procedure similar to as-isolated spectra) followed by drying under a tent atmosphere without an external gas stream.

#### Molecular dynamics simulation details

All molecular dynamics (MD) simulations were conducted using the GROMACS software package, version 2021.1.[12] The initial structure for the wild-type (WT) ToHyd [FeFe]-hydrogenase was derived from an AlphaFold prediction model (ColabFold v1.3.0, AlphaFold2 using MMseqs2)[13,14] and refined using the MolProbity webtool.[15] The H-cluster and accessory FeS clusters were incorporated into the prediction model by structural superimposition with the X-ray crystal structure of CpI (PDB ID: 4XDC).[16] The iP207_dC211 and I208F ToHyd [FeFe]-

hydrogenase variants were generated from the WT protein structure through in-silico mutation. Specifically, the proton-transporting cysteine-containing loop in the iP207_dC211 variant was constructed by mimicking the corresponding loop from CpI.

The CHARMM36 protein force field and the CHARMM-specific TIP3P water model were utilized for the simulations.[17] This study focused on the $H_{ox}$ state of ToHyd [FeFe]-hydrogenases. Force field parameters for the H-cluster and accessory FeS clusters, including the coordinating cysteine and histidine residues, were adopted from the work of Chang et al.,[18] with additional modifications as recommended by McCullagh and Voth.[19]

The predicted protein structures initially lacked buried water molecules. These were added using the WarPP webtool.[20] After placing the buried water molecules, each protein was fully solvated with 16519 water molecules and the net charge of the systems was neutralized by adding 17 $Na^+$ ions. The simulations were carried out with periodic boundary conditions in dodecahedron-shaped simulation boxes, encompassing a total of 56208 atoms for the wild-type, 56211 atoms for the iP207_dC211 variant, and 56209 atoms for the I208F variant.

Before the MD simulations, the systems underwent energy minimization using 10,000 steps of steepest descent. This was followed by stepwise equilibration with harmonic position restraints applied to various atom sets. The equilibration phase began with an NVT simulation, where the system temperature was increased from 0 K to 300 K over 0.2 ns. During this NVT simulation, position restraints with force constants of 1,000 kJ/mol/nm$^2$ were applied to all non-hydrogen atoms of the protein and FeS clusters. The equilibration continued for an additional 2.5 ns in the NPT ensemble. During the first 0.5 ns of the NPT equilibration, the same position restraints were maintained on all non-hydrogen atoms of the protein and FeS clusters. In the subsequent 2.0 ns of NPT equilibration, restraints were applied only to the protein backbone atoms, allowing the protein side chains, FeS clusters, and surrounding water molecules to relax. Finally, three 1000 ns production simulations were conducted within the NPT ensemble at 300 K for each of the three protein systems. These simulations used the equilibrated systems as starting conditions, with different random seeds to generate the initial atomic velocities from a Maxwell-Boltzmann distribution. For analysis, the last 500 ns of each simulation were utilized. The identical MD simulation protocols were successfully employed in previous studies with CpI and CbA5H.[6,21] The analyses included distance distribution between the $Fe_d$ atom and the thiol S-atom of capping cysteine (Cys) residue, hydrogen bond analysis, and hydrophobic cluster analysis. Hydrophobic cluster formations composed of sidechains from isoleucine (Ile), leucine (Leu), and valine (Val) residues were analyzed using the Protein Tools web server.[22]

During the MD simulations, the temperature was maintained at 300 K using the velocity rescaling thermostat by Bussi and coworkers,[23] with a coupling time constant of 0.1 ps. A constant pressure of 1 bar was maintained using an isotropic weak coupling Berendsen barostat,[24] with a coupling time constant of 2 ps and a compressibility of $4.5 \times 10^{-5}$ bar$^{-1}$. Short-range Coulomb and Lennard-Jones 6,12 interactions were computed using a buffered Verlet pair list,[25] with potentials smoothly transitioning to zero at a 1.2 nm cutoff and forces smoothly shifted to zero between 1.0 and 1.2 nm. Long-range electrostatic interactions were treated with the particle mesh Ewald (PME) method, using a 0.12 nm grid spacing.[26] The LINCS algorithm was used to constrain all protein bonds involving hydrogen atoms,[27] while the SETTLE algorithm was employed to constrain all internal degrees of freedom of water molecules.[28] This setup allowed for the integration of motion equations with 2 fs time steps.

### Supporting figures and tables

#### Figure S1. Comparison of proton transporting cysteine bearing loop of group A and group B hydrogenases

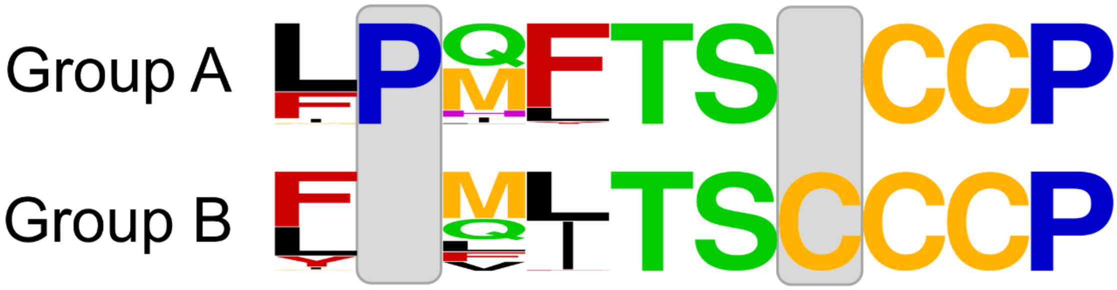


**Figure S1:** The WebLogo[29,30] representation of the sequence alignment for 144 [FeFe]-hydrogenases was generated using Clustal Omega.[31] The alignment includes all 71 Group B hydrogenase sequences available on HydDB[1] (featuring the TSCCCP motif) along with the CpIII sequence. Since the most characterized Group A hydrogenases belong to Group A1, the alignment includes 68 Group A1 sequences with the TSCCP motif (out of 180 Group A1 sequences available in HydDB)[1], along with CpI, CbA5H, CrHydA1, and DdH sequences. The additional cysteine in Group B and the conserved proline in Group A are highlighted with grey boxes. The Clustal Omega sequence alignment of 144 [FeFe]-hydrogenases is uploaded as a separate pdf file.

#### Figure S2. SDS PAGE of ToHydA WT and variants after purification

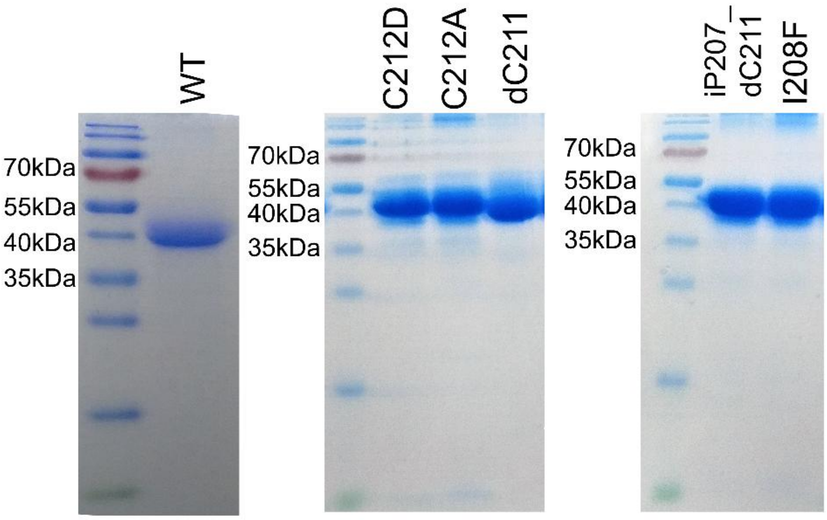


**Figure S2:** In left panel, protein marker and ToHydA WT; In middle panel, protein marker and variants C212D, C212A and dC211; and in right panel, protein marker and variants iP207_dC211 and I208F.

#### Figure S3. Fe content of ToHydA WT

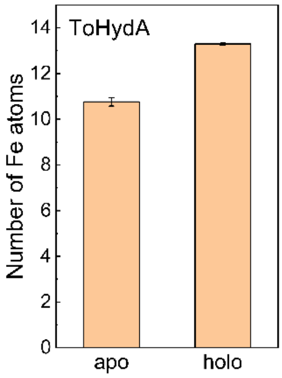


**Figure S3:** The iron content (Fe atoms per protein molecule) of ToHydA WT was determined using inductively coupled plasma-optical emission spectroscopy (ICP-OES) for both the holo form (with $[2Fe]_H$) and the apo form (without $[2Fe]_H$).

#### Figure S4. *In-vitro* $H_2$ production activity of ToHydA WT and variants at elevated temperatures

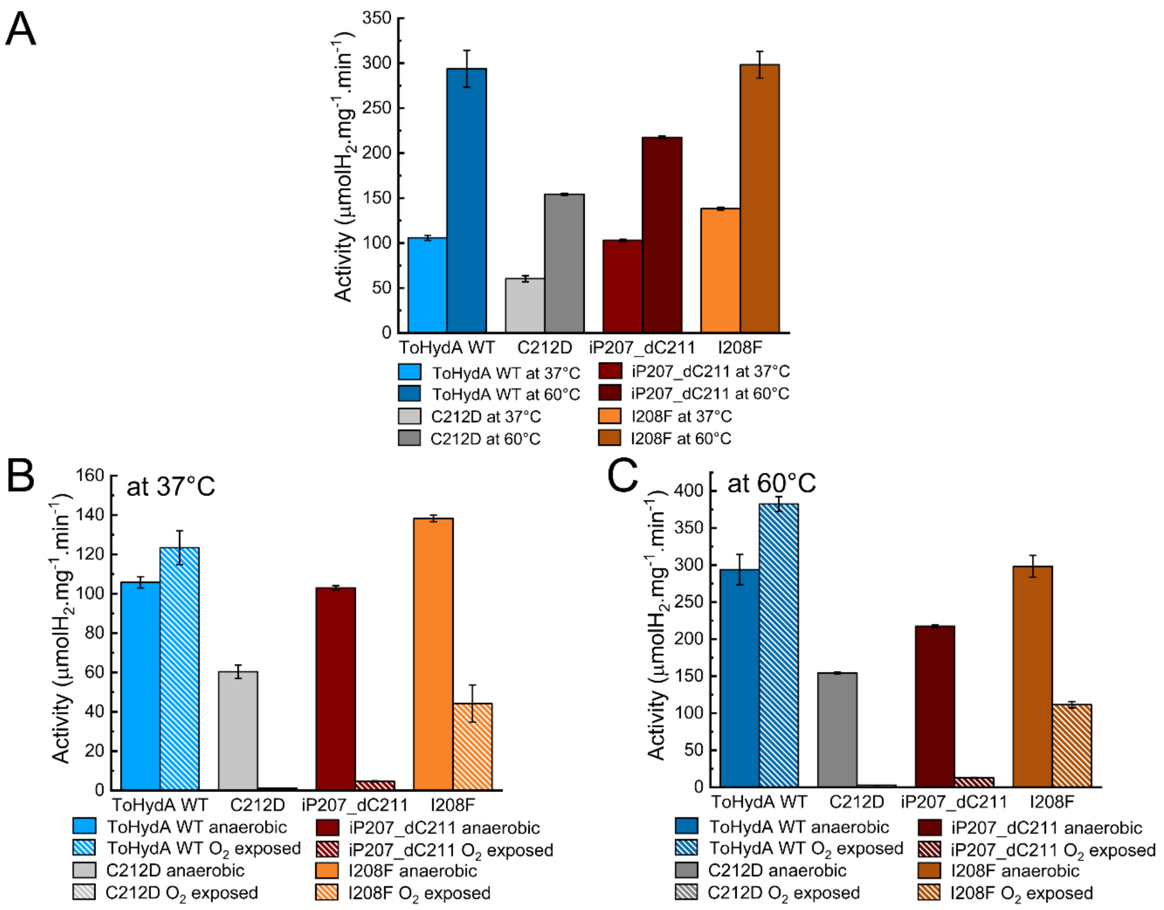


**Figure S4:** A. In-vitro $H_2$ production activity of ToHydA WT and variants (anaerobic proteins) measured at 37°C and 60°C. B. In-vitro $H_2$ production activity of ToHydA WT and variants before and after 20 minutes of $O_2$ incubation at 37°C, with $H_2$ production measured at 37°C. C. In-vitro $H_2$ production activity of ToHydA WT and variants before and after 20 minutes of $O_2$ incubation at 60°C, with $H_2$ production measured at 60°C. For $O_2$ incubation, 6 µL of 25-30 mg/mL protein samples in 100 mM Tris-HCl buffer (pH8) containing 2mM NaDT were exposed to air at elevated temperatures (37°C or 60°C).

#### Figure S5. Electrochemistry

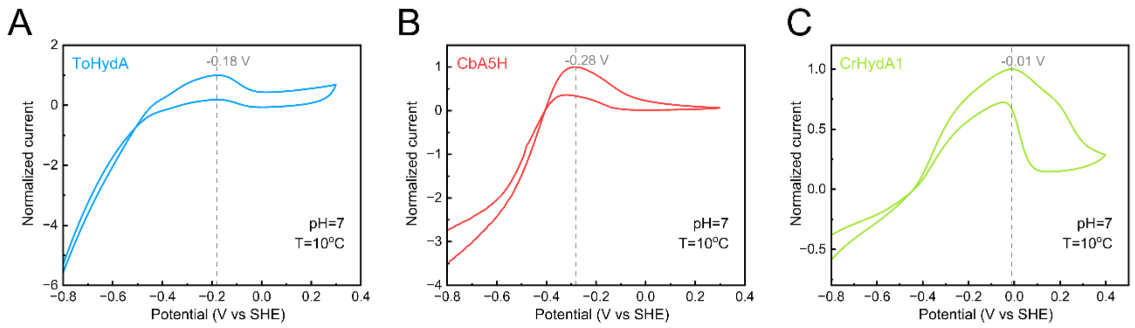


**Figure S5:** Cyclic voltammograms of A. ToHydA WT B. CbA5H WT and C. CrHydA1 WT (in 100mM potassium phosphate buffer with 10mM NaCl, temperature of 10 °C, at pH 7, 1 atm. of $H_2$, scan rate of 0.5 mV/s, electrode rotation rate of 3000 rpm). The currents are normalized by dividing with maximum oxidation current. The potential at which oxidative inactivation occurs through cysteine binding at $Fe_d$ in ToHydA is distinct from both

CbA5H and CrHydA1. However, in the case of CrHydA1, the same occurs through a Cl- ion binding at Fed site. Due to the low current observed in ToHydA WT, the measurements without the addition of NaCl were not feasible.

#### Figure S6. FTIR spectra of as-isolated ToHydA WT with 2mM NaDT and without NaDT.

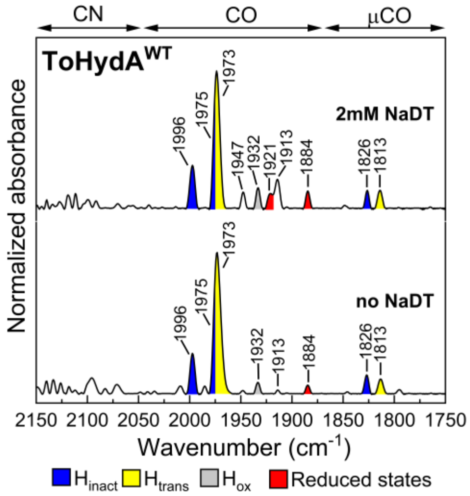


**Figure S6:** FTIR spectra of as-isolated ToHydA WT with 2mM NaDT and without NaDT. The spectra of the proteins are normalized to the second amide band (1535-1545 cm$^{-1}$). 0.4 - 0.5 mM protein sample was prepared in 100 mM Tris/HCl buffer (pH 8.0).

#### Figure S7. The "$H_2$–$N_2$" ("$D_2$-$N_2$" in case of $D_2O$) difference FTIR spectrum of ToHydA in $H_2O$ and $D_2O$

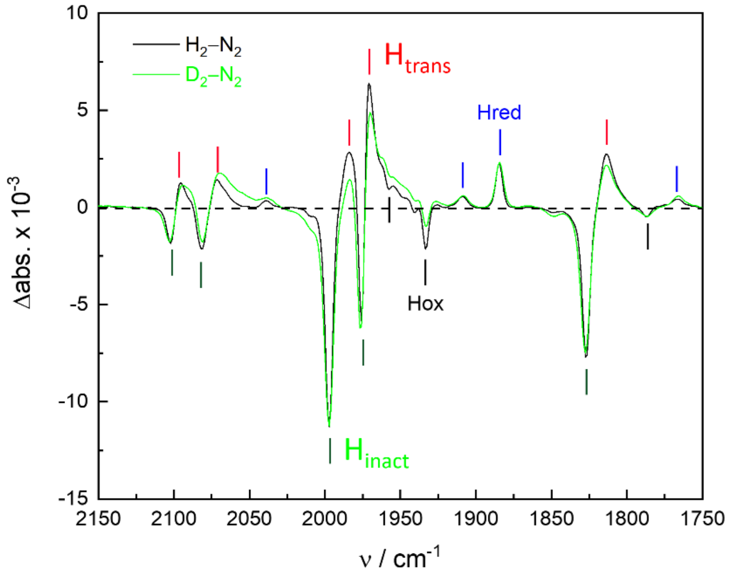


**Figure S7*:*** The "$H_2$–$N_2$" ("$D_2$-$N_2$" in case of $D_2O$) difference FTIR spectrum of ToHydA in $H_2O$ and $D_2O$.

Figure S8. *In-vitro* $H_2$ production activity of ToHydA$^{WT}$, CbA5H$^{WT}$, and CpI$^{WT}$

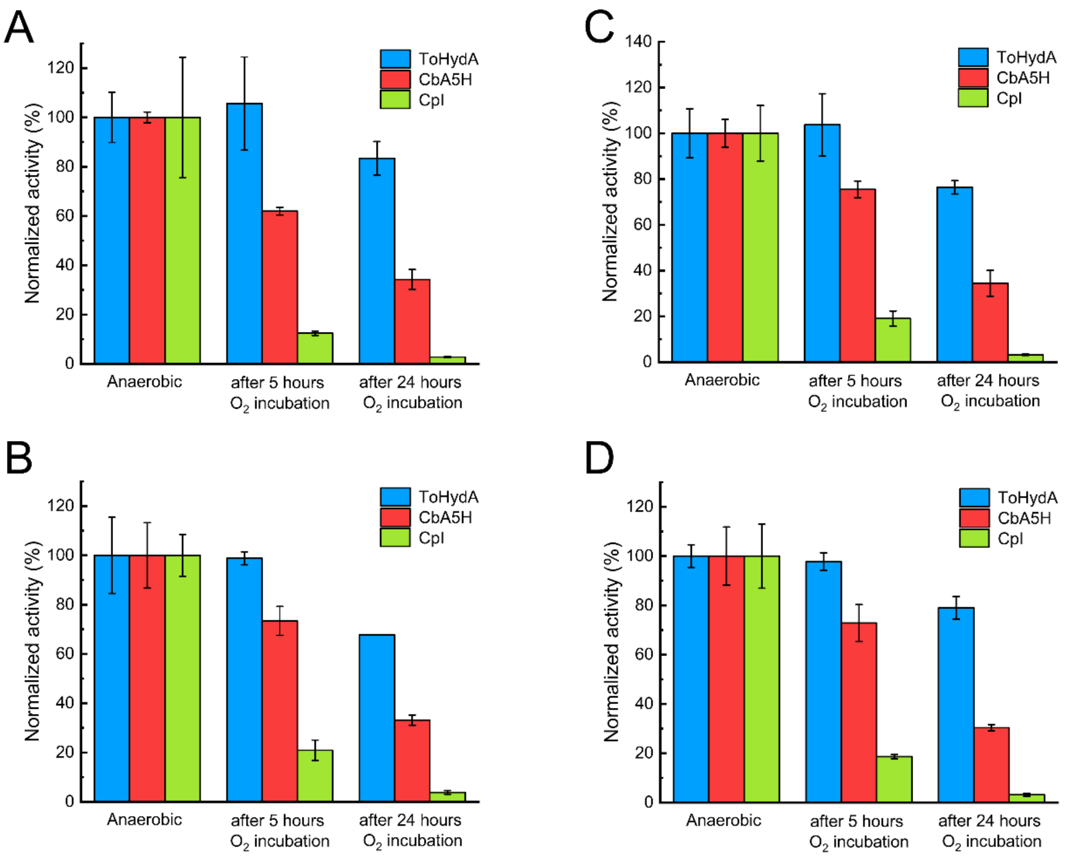


**Figure S8:** *In-vitro* methyl viologen (MV) mediated $H_2$ production activities of ToHydA, CbA5H and CpI anaerobic and after 5h and 24h $O_2$ incubation. A. and B. represent technical replicates from a single protein isolation (Biological replicate 1) measured on different days. C. and D. are technical replicates from a separate protein isolation (Biological replicate 2), also measured on different days. Each dataset is presented as the average of three measurements, along with the standard deviation. For $O_2$ incubation, 10 µL of 15-20 mg/mL protein samples in 100 mM Tris-HCl buffer (pH8) containing 2mM NaDT were stored under air for 5 and 24 hours at 8°C. $H_2$ production was measured at 37°C.

Figure S9: UV-Vis spectra of ToHydA WT and its variants.

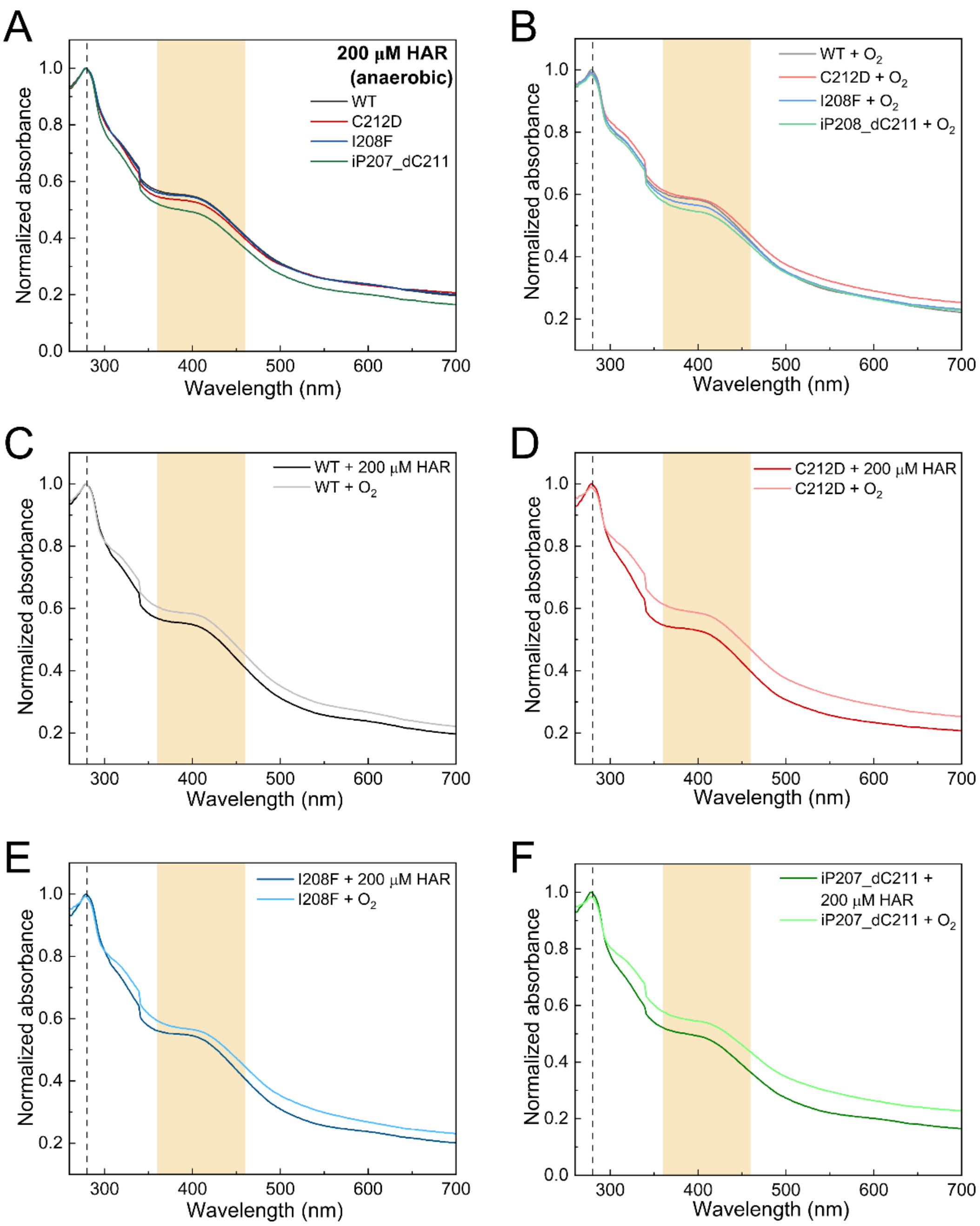


**Figure S9: UV-Vis spectra of ToHydA WT and its variants.** A. UV-Vis spectra of anaerobically oxidized ToHydA WT and variants, obtained by adding 200 µM HAR to 10 µM protein sample. B. UV-Vis spectra of $O_2$-treated ToHydA WT and variants. 10 µM protein samples were incubated with air for 20 minutes. C-F. Comparison of UV-Vis spectra between anaerobically and aerobically oxidized samples. All spectra are normalized to the 280 nm peak (marked by black dashed line), which corresponds to the absorption of aromatic amino acids in the protein samples. The yellow-shaded region spanning 360–460 nm indicates the characteristic absorption range of [4Fe-4S] clusters. The spectra were recorded at 25°C in 0.1 mM Tris-HCl buffer (pH 8.0). Before the measurements, NaDT was removed by buffer exchange.

Figure S10. Sequence of the C212-bearing loop for ToHydA WT and variants

| | | | | | | | | | | |
|---|---|---|---|---|---|---|---|---|---|---|
| ToHydA$^{WT}$ | $F_{206}$ | – | $M_{207}$ | $I_{208}$ | $T_{209}$ | $S_{210}$ | $C_{211}$ | $C_{212}$ | $C_{213}$ | $P_{214}$ |
| ToHydA$^{C212D}$ | $F_{206}$ | – | $M_{207}$ | $I_{208}$ | $T_{209}$ | $S_{210}$ | $C_{211}$ | **$D_{212}$** | $C_{213}$ | $P_{214}$ |
| ToHydA$^{C212A}$ | $F_{206}$ | – | $M_{207}$ | $I_{208}$ | $T_{209}$ | $S_{210}$ | $C_{211}$ | **$A_{212}$** | $C_{213}$ | $P_{214}$ |
| ToHydA$^{dC211}$ | $F_{206}$ | – | $M_{207}$ | $I_{208}$ | $T_{209}$ | $S_{210}$ | **–** | $C_{211}$ | $C_{212}$ | $P_{213}$ |
| ToHydA$^{iP207_dC211}$ | $F_{206}$ | **$P_{207}$** | $M_{208}$ | $I_{209}$ | $T_{210}$ | $S_{211}$ | **–** | $C_{212}$ | $C_{213}$ | $P_{214}$ |
| ToHydA$^{I208F}$ | $F_{206}$ | – | $M_{207}$ | **$F_{208}$** | $T_{209}$ | $S_{210}$ | $C_{211}$ | $C_{212}$ | $C_{213}$ | $P_{214}$ |

**Figure S10:** Sequence of the C212-bearing loop for ToHydA WT and variants studied in this work. The point mutation sites are highlighted in the figure.

Figure S11. *In-vitro* $H_2$ production activity of ToHydA WT and variants

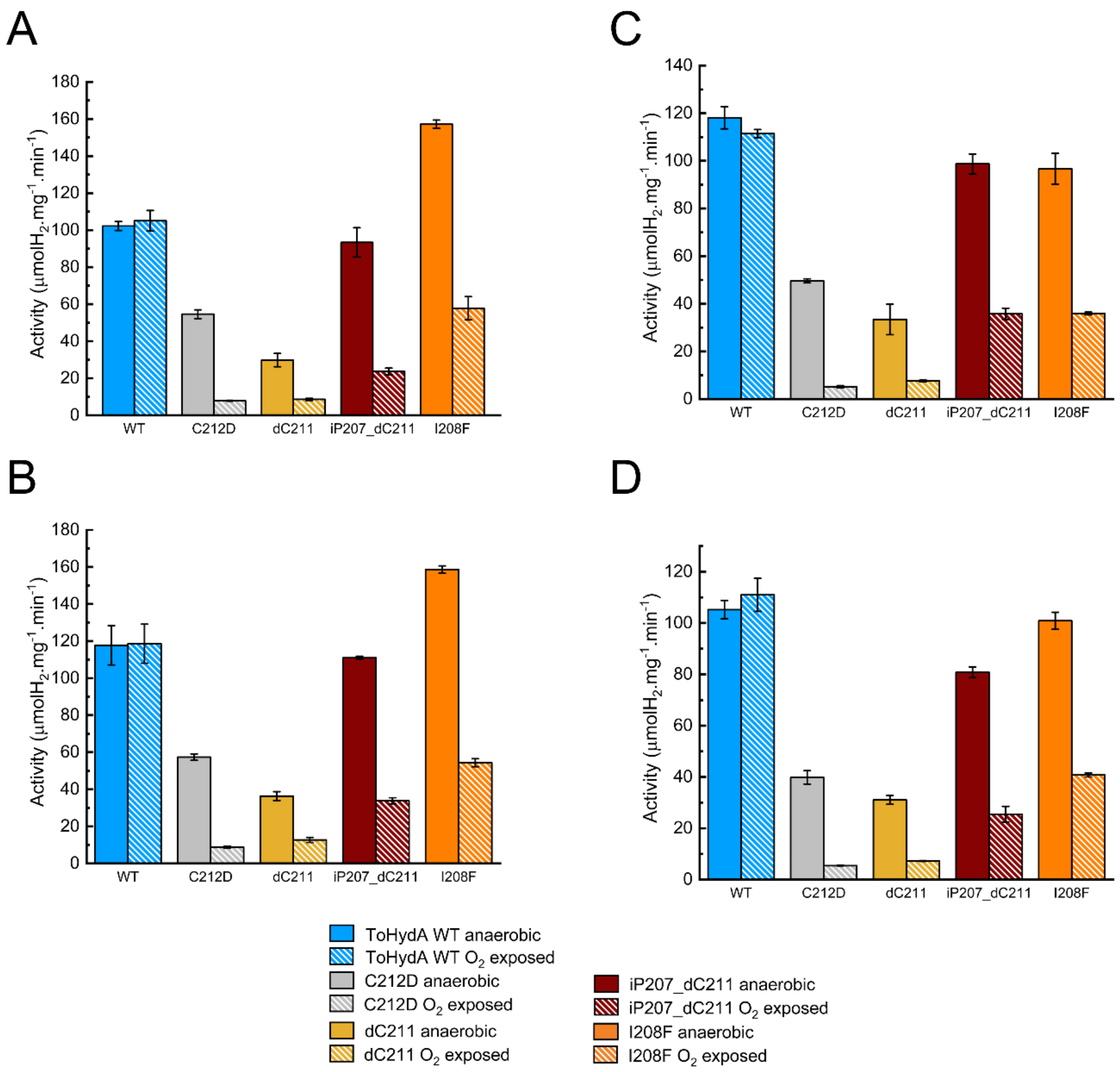


**Figure S11:** *In-vitro* MV-mediated $H_2$ production activity of ToHydA WT and variants before and after 20 minutes $O_2$ incubation. A. and B. represent technical replicates from a single protein isolation (Biological replicate 1) measured on different days. C. and D. are technical replicates from a separate protein isolation (Biological replicate 2), also measured on different days. Each dataset is presented as the average of three measurements, along with the standard deviation. For $O_2$ incubation, 6 µL of 25-30 mg/mL protein samples in 100 mM Tris-HCl buffer (pH8) containing 2mM NaDT were exposed to air on ice (at 4°C) for 20 minutes. $H_2$ production was measured at 37°C.

Figure S12. Biochemical and ATR-FTIR spectroscopic characterization of ToHydA dC211

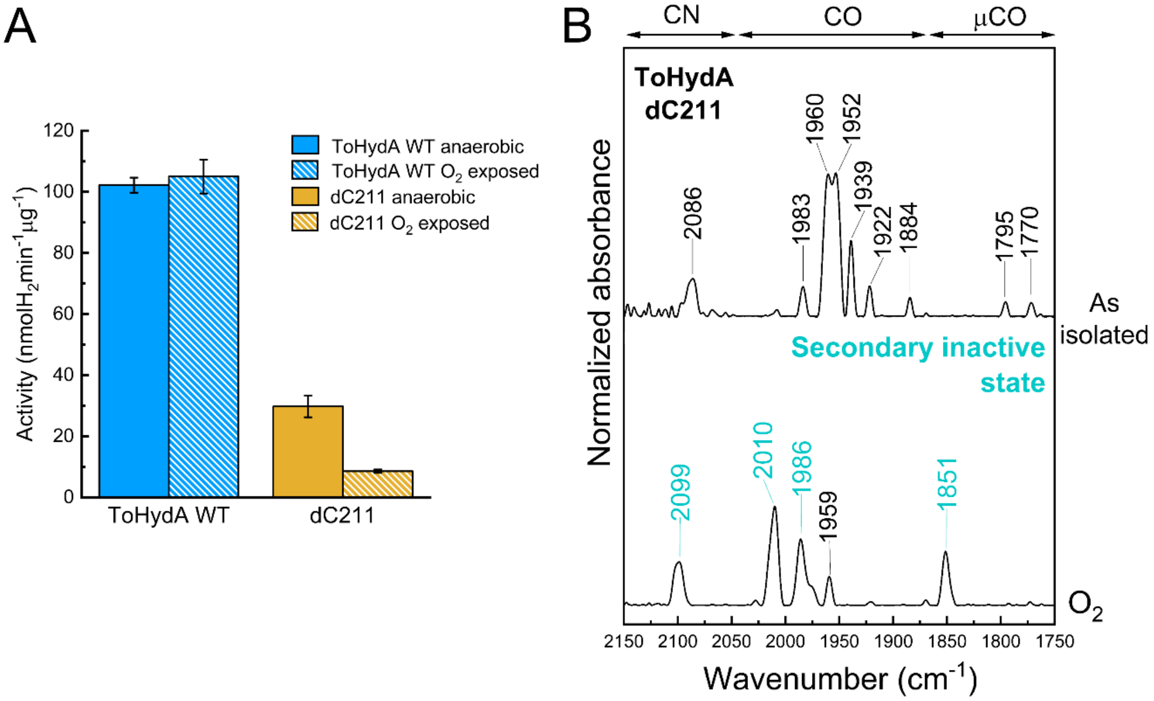


**Figure S12:** A. *In-vitro* $H_2$ production activity of ToHydA WT and dC211 variant before and after 20 minutes $O_2$ incubation. For $O_2$ incubation, 6 µL of 25-30 mg/mL protein samples in 100 mM Tris-HCl buffer (pH8) containing 2mM NaDT were exposed to air on ice (at 4°C) for 20 minutes. $H_2$ production was measured at 37°C. B. FTIR spectra of ToHydA dC211 variant in the as-isolated state and after $O_2$ exposure. Gas purging experiment started from as-isolated state. The spectra of the proteins are normalized to the second amide band (1535-1545 $cm^{-1}$). 0.4 - 0.5 mM protein sample was prepared in 100 mM Tris/HCl buffer (pH 8.0) containing 2 mM NaDT.

#### Figure S13. ATR-FTIR spectra of variants ToHydA C212A and ToHydA iP207_dC211

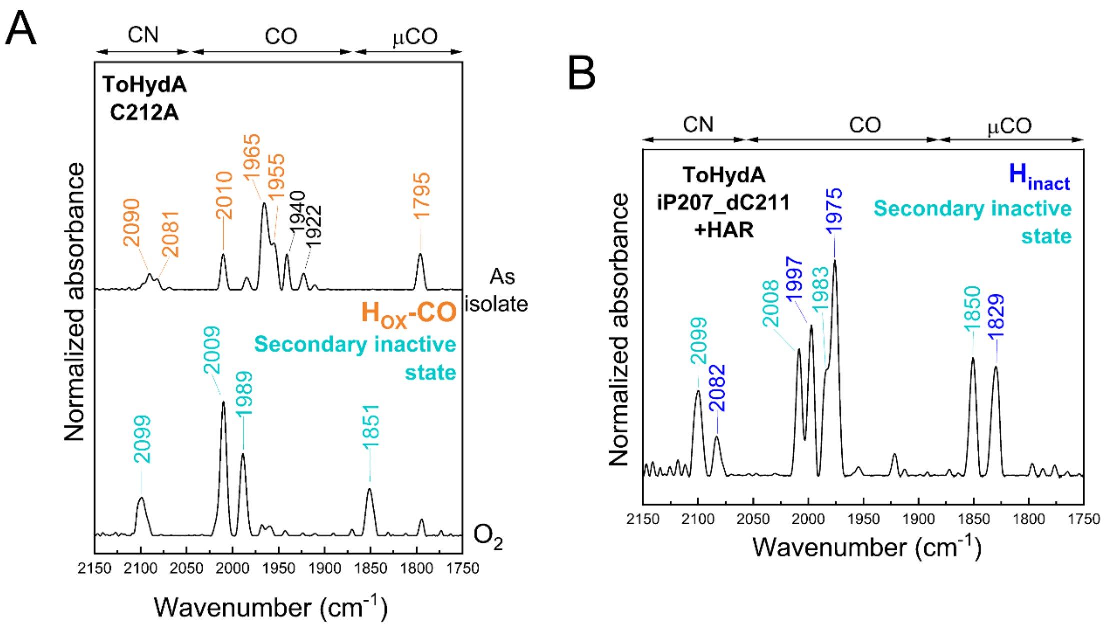


**Figure S13:** A. FTIR spectra of variant ToHydA C212A (mutation site shown in Figure 1 and SI Figure S10; primers used for QC-PCR are listed in Table 1; SDS-PAGE shown in SI Figure S2) in the as-isolated state and after $O_2$ exposure. B. The FTIR spectrum of variant ToHydA iP207_dC211 in the presence of 100mM HAR. HAR possesses a midpoint potential of +55 mV is therefore an oxidant for [FeFe]-hydrogenases. Gas purging experiment started from as-isolated state. The spectra of the proteins are normalized to the second amide band (1535-1545 cm$^{-1}$). 0.4 - 0.5 mM protein sample was prepared in 100 mM Tris/HCl buffer (pH 8.0) containing 2 mM NaDT. Overall, the spectrum appears to be comprised of two different inactive states. The $H_{inact}$ state (with blue labellings) is very similar to that observed in WT protein. The secondary inactive state (colored cyan) resembles a $H_2O/OH^-$ bound state which was identified in Group D [FeFe]-hydrogenase and proposed to bind a water or hydroxide ion at the H-cluster.[32]

#### Figure S14. Sequence Comparison of LPA motif of CbA5H with ToHydA and CpI

| | | | | | | | | | | | | | | | | |
|---|---|---|---|---|---|---|---|---|---|---|---|---|---|---|---|---|
| **ToHydA** | M | $I_{208}$ | T | S | C | $C_{212}$ | C | P | ... | M | $V_{231}$ | S | ... | N | $F_{365}$ | V |
| **CbA5H** | I | **$L_{364}$** | T | S | - | $C_{367}$ | C | P | ... | V | **$P_{386}$** | S | ... | H | **$A_{561}$** | I |
| **CpI** | M | $F_{296}$ | T | S | - | $C_{299}$ | C | P | ... | N | $L_{364}$ | S | ... | H | $F_{493}$ | I |

**Figure S14:** A sequence alignment comparing the LPA (comprised of L364, P386 and A561) motif of CbA5H with ToHydA and CpI. L364 of CbA5H is homologous to I208 of ToHydA and F296 of CpI, as highlighted in green. P386 and A561 of CbA5H are homologous to V231 and F365 of ToHydA, and L364 and F493 of CpI, respectively, as highlighted in orange and grey. The proton transporting cysteine is also indicated in blue. As isoleucine and leucine are structural isomers, the LPA motif is partially conserved in ToHydA. Conversely, the other two homologous positions in ToHydA are different from CbA5H, however closely identical to CpI, as leucine and valine are functionally similar amino acids.

#### Figure S15. ToHydA WT whole protein representation

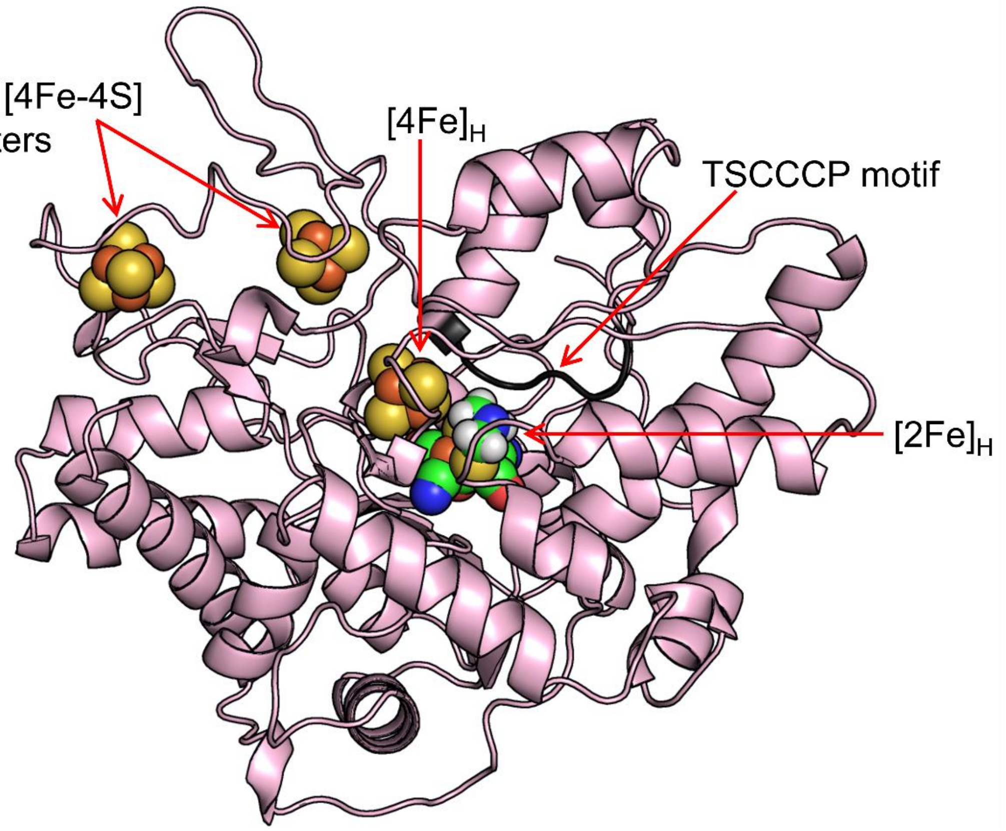


**Figure S15:** A representative MD-snapshot showing whole protein scaffold (in cartoon representation) of ToHydA WT with integrated FeS clusters (in sphere representation). The TSCCCP motif is highlighted in dark grey. The structure of ToHydA WT used for rendering the figure is uploaded as a separate pdb file.

#### Figure S16. Hydrophobic cluster formation in CbA5H, CrHydA1 and CpI

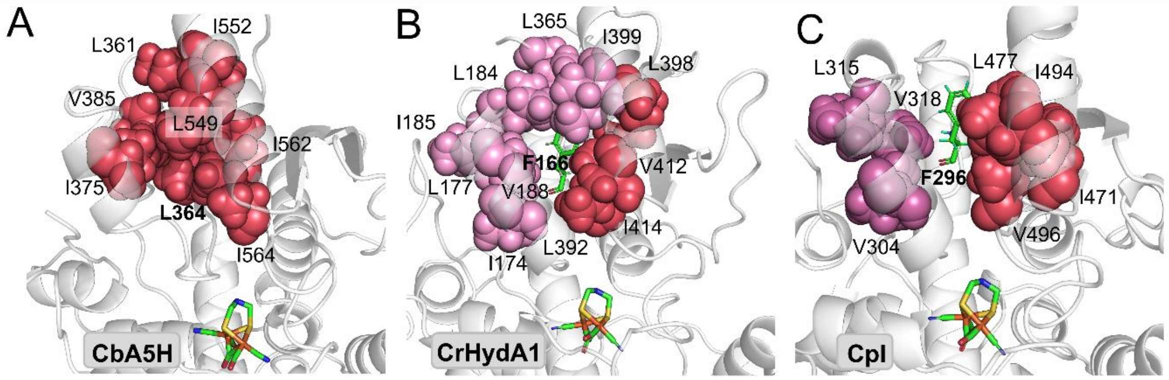


**Figure S16:** The hydrophobic cluster formation centered around the proton transporting cysteine bearing loop, involving Val (V), Ile (I), and Leu (L) residues are illustrated as van der Waals spheres for group A hydrogenases. A. The hydrophobic cluster is preserved in CbA5H as found in the case of ToHydA WT and iP207_dC211. B and C. In CrHydA1 and CpI, this cluster splits into two smaller clusters due to phenylalanine, similar to ToHydA I208F variant. The equilibrated structures of CpI and CbA5H taken from our previous works were used for probing the hydrophobic cluster formation,[6,21] while x-ray crystal structure of CrHydA1 (PBD ID 3LX4) was used for this analysis.[33]

Figure S17. probability of hydrogen bond formation between the C212-bearing loop and neighboring loops

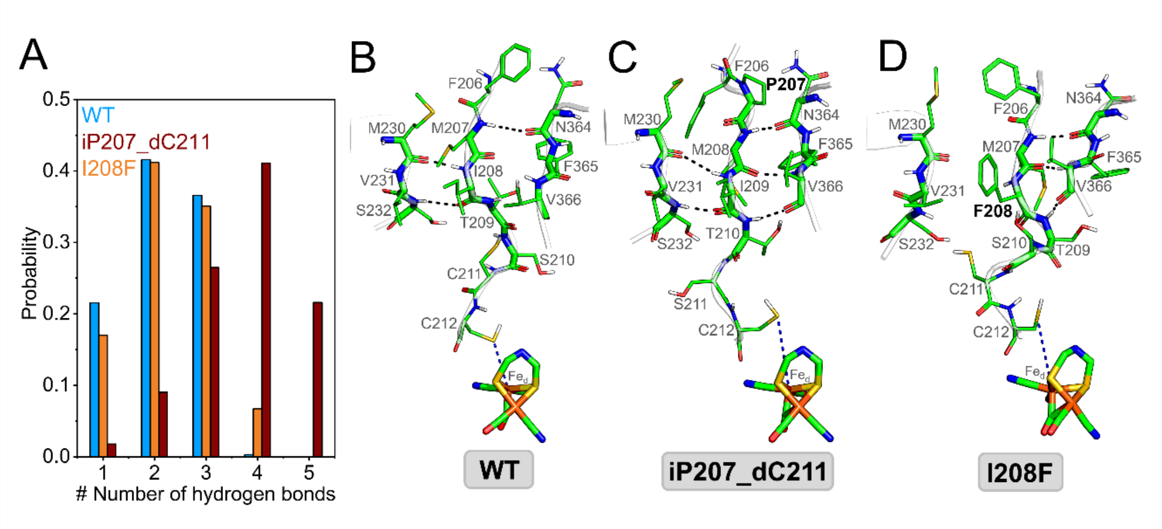


**Figure S17:** The probability of hydrogen bond formation between the C212-bearing loop and neighboring loops in ToHydA WT and its variants, along with representative MD snapshots illustrating the hydrogen bond network.

Figure S18: Proposed model for $H_{inact}$ formation in ToHydA

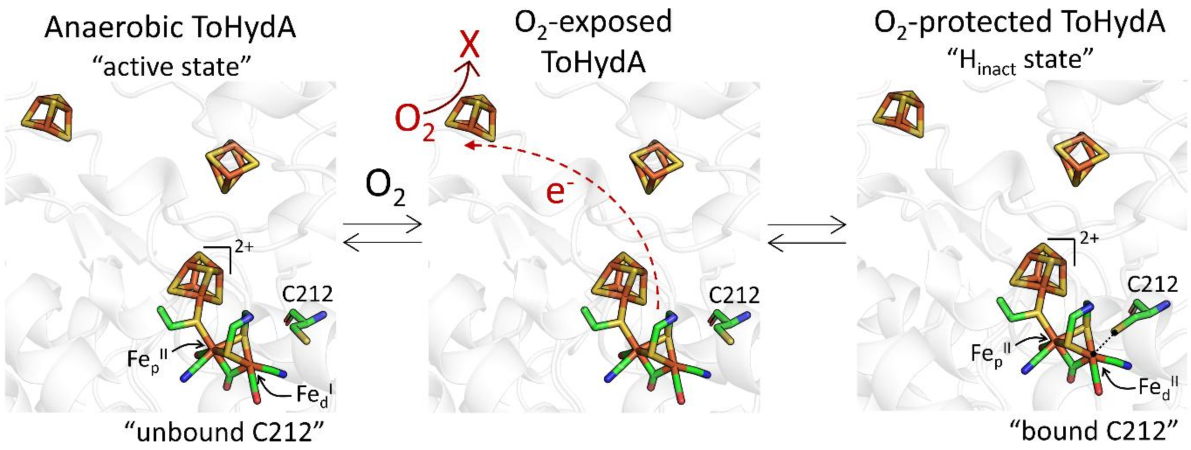


**Figure S18:** Proposed model for $H_{inact}$ formation in ToHydA. Under anaerobic conditions, C212 remains unbound. Upon $O_2$ exposure, the accessory [4Fe-4S] clusters and the H-cluster undergo oxidation and $O_2$ is reduced (the reduced species are denoted as X; X could $O_2^-$, $O_2^{2-}$ etc. depending on the number of electrons available). . This is followed by the binding of C212 to the oxidized $Fe_d$, leading to the formation of the $H_{inact}$ state.

Figure S19. Water molecules near active site of ToHydA iP207_dC211 in MD simulation

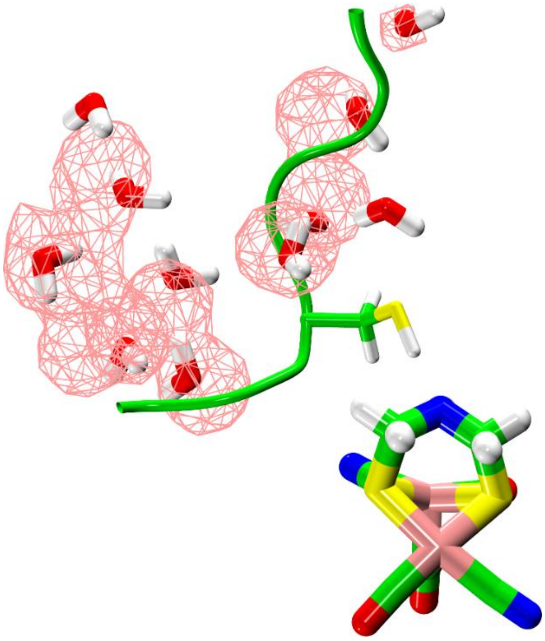

**Figure S19:** Active site water molecules around the H-cluster of $H_{inact}$-impaired iP207_dC211 variant. The pink mesh indicates the averaged water occupancy around the H-cluster throughout the MD simulation. The threshold of the water occupancy map shown in the figure is set to 30%.

Table S1. Generation of site-directed mutagenesis variants of ToHydA. Primers used for QuikChange PCR (QC-PCR) to generate variants of ToHydA.

| | |
|---|---|
| **C212D_for:** | CCAGCTGTGATTGTCCGCCTTTTGTTCG |
| **C212D_rev:** | GGCGGACAATCACAGCTGGTAATCATGAATTTATCG |
| **C212A_for:** | CCAGCTGTGCGTGTCCGCCTTTTGTTCG |
| **C212A_rev:** | GGCGGACACGCACAGCTGGTAATCATGAATTTATCG |
| **dC211_for:** | TTACCAGCTGTTGTCCGCCTTTTGTTCG |
| **dC211_rev:** | GACAACAGCTGGTAATCATGAATTTATCGCC |
| **iP207_dC211_for:** | TAAATTCCCGATGATTACCAGCTGTTGTCCGC |
| **iP207_dC211_rev:** | ATCATCGGGAATTTATCGCCACGATCCATACG |
| **I208F_for:** | ATTCATGTTTACCAGCTGTTGTTGTCCGC |
| **I208F_rev:** | CTGGTAAACATGAATTTATCGCCACGATCC |
